# Category learning drives cortical attentional gain to diagnostic dimensions

**DOI:** 10.1101/2024.09.21.614258

**Authors:** Sahil Luthra, Raha N. Razin, Adam T. Tierney, Lori L. Holt, Frederic Dick

## Abstract

Humans and other animals develop remarkable perceptual and cognitive specializations for identifying, differentiating, and acting on classes of ecologically important signals. This expertise is flexible enough to support diverse perceptual judgments: a voice, for example, simultaneously conveys what a talker says, as well as myriad cues about her identity and state. Expert perception across complex signals thus involves discovering and learning regularities that best inform diverse perceptual judgments, as well as weighting this information flexibly as task demands change. Here, we test whether this flexibility may involve endogenous attentional gain. We use two prospective auditory category learning tasks to relate a complex, entirely novel soundscape to four classes of “alien identity” and two classes of “alien size.” Identity, but not size, categorization requires discovery and learning of patterned acoustic input situated in one of two simultaneous, non-overlapping frequency bands. This allows us to capitalize on the coarsely segregated frequency-band-specific channels tiling auditory cortex, using fMRI to ask whether category-relevant perceptual information present in one frequency band is prioritized relative to simultaneous, uninformative information in the other frequency band. Among participants expert at alien identity categorization, we observe prioritization of the identity-diagnostic frequency band that persists even when the diagnostic information becomes irrelevant in the size categorization task. Tellingly, the neural selectivity evoked implicitly in the identity categorization task aligns with that in an independent task, where activation is driven by explicit and sustained selective attention to pure tones in one or the other frequency band. Additionally, the learning trajectories taken to achieve expert-level categorization leave fingerprints on the patterns of neural activity associated with the diagnostic dimension. In all, this indicates that acquiring categories can drive the emergence of acquired attentional gain to category-diagnostic input dimensions.

---

Learners of all ages discover structured perceptual input to achieve their goals, and even simple sights and sounds convey rich information that can flexibly support multiple behaviors. A single utterance simultaneously conveys a talker’s age (Lavan, 2022) and socioeconomic class (Kraus et al. 2019), whether she is a stranger or a loved one (Holmes et al., 2018), and if she is requesting her *bill* or her *pill* (Murphy et al. 2024). A glimpse of her face communicates her trustworthiness (Todorov et al. 2009), emotion (Krumhuber et al. 2023), and identity (Young et al. 2020). Acoustic characteristics of her footsteps can convey her gender (Li et al., 1991), identity (Algermissen & Hörnlein 2021), health, and weight (Cunningham & McGregor, 2024). These underlying patterns are carried across multiple perceptual dimensions and – especially in the case of sound – often require integration of information across time. Successful perception thus requires that we discover and learn distinct input patterns that best inform specific judgments. Flexibility in weighting the most diagnostic patterns of perceptual cues or dimensions for the task at hand is crucial for effective face and voice identification, object recognition, and at least in humans, speech comprehension.

Prominent theories of categorization have posited that learning may drive endogenous attention to be directed toward diagnostic perceptual dimensions in a task-dependent manner (e.g., Gao et al. 2024; Nosofsky, 1986). These learned attentional biases are proposed to weight perceptual dimensions most relevant to oft-performed tasks and behaviors (Van Gulick & Gauthier, 2014; De Baene et al. 2008; O’Bryan et al. 2024), and to language-specific cues (Chandrasekaran, Sampath, & Wong, 2010; Jasmin, Sun, & Tierney, 2021; Petrova et al., 2023). For complex skills like spoken language and object recognition, it can take years or even decades to learn which perceptual dimensions are informative for a given behavioral goal and to optimally weight or attend to them (e.g., Creel et al., 2023; Idemaru & Holt, 2013; McMurray et al. 2018; Yurovsky et al. 2013; Yurovsky & Frank, 2017). Learners also progress along different trajectories to reach equivalent performance outcomes (Roark et al. 2024; Reetzke et al. 2018). Indeed, distinct perceptual and neural strategies are reflected in developmental neuroimaging studies that demonstrate that there can be quite different weightings of neural activation across brain regions, even among children or adults who perform similarly on a task (e.g., Brown et al. 2005; Stiles et al. 2003; Karmiloff-Smith, 2010; Krishnan et al., 2014; Yates et al., 2021). In auditory categorization tasks, both mouse (Colling et al., 2025) and human (Sheth et al., 2025; Roark & Chandrasekaran, 2023) learners exhibit idiosyncratic patterns of behavior over short-term training. These individual differences may in turn reflect where participants are positioned in a longer-term learning trajectory, over which diagnostic input dimensions are discovered and tuned through experience.

Previous functional MRI work has shown that short-term category training can shape distributed auditory (Ley et al., 2012) and visual (Folstein et al., 2014) cortical representations. Here, we establish whether learning a new behavior across an informationally rich perceptual space a) drives distinct patterns of cortical activity that are spatially predictable given the cortical mapping of the category-diagnostic dimension, and b) does so in a way that is consistent with acquired endogenous attention to the perceptual dimensions most diagnostic to the task. By examining cohorts of learners who reach equivalent levels of expertise in this new domain, we also ask whether patterns of cortical activity reflect the distinct trajectories learners take to achieve expertise. To do so, we adopt a prospective learning approach in which naïve adults learn to categorize entirely novel sounds drawn from a complex, multidimensional perceptual space composed of over 36,000 sounds that vary across complex, not-readily-verbalizable acoustic dimensions (Obasih et al., 2023). Both the sound exemplars and the categories we have defined across them are entirely novel to listeners, as are the ‘space alien identity’ and ‘space alien size’ visual cues.

The space alien sounds possess information in two simultaneous, non-overlapping frequency bands (**Fig 1**), with each band having highly similar and complex acoustic information. Each of four aliens’ identities is conveyed by spectrotemporal patterns within one or the other of these frequency bands. This allows us to take advantage of the tonotopic mapping of auditory cortex, where we expect acoustic information in each frequency band to be preferentially processed in different spatial subregions of the multiple tonotopic maps that lie across the superior temporal lobe.

**Figure 1.**
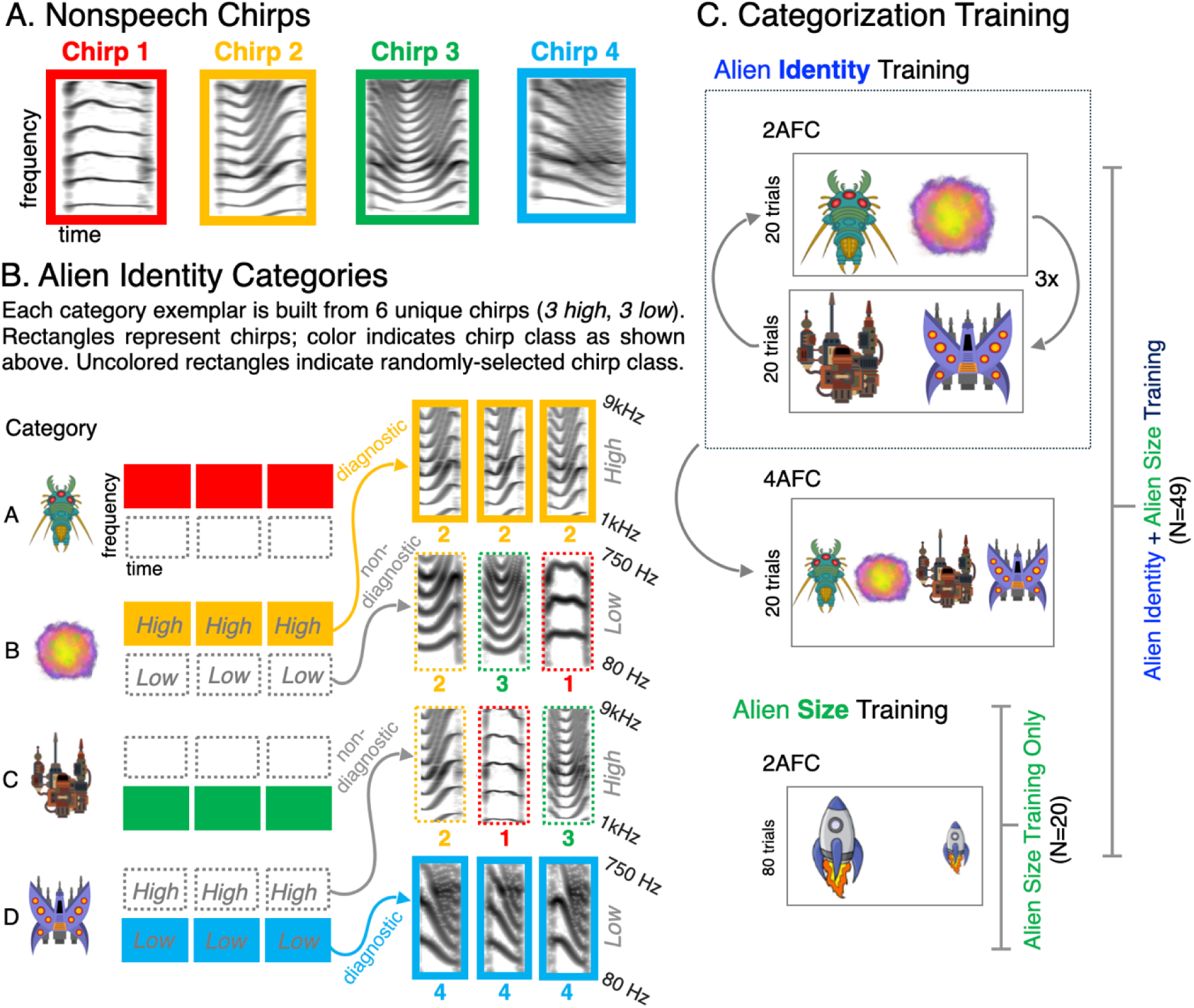
Stimuli. **(A) Nonspeech Chirps.** We extracted the fundamental frequency (F0) contour from natural speech recordings of single-syllable words varying in Mandarin lexical tone contour across four (2 female) native Mandarin speakers. This yielded 80 acoustically unique chirps from each of the four classes of Mandarin lexical tone contour (roughly: level, rising, dipping or falling; see Obasih et al. 2023). Spectrograms show one representative chirp from each class (indicated by the colored outline). **(B) Alien Identity Categories.** We next shifted each nonspeech chirp into two non-overlapping frequency bands (low: ∼80-750 Hz, high: ∼1000-9000 Hz), creating two sets of chirps to serve as building blocks for creating novel auditory category exemplars. For each category, one frequency band conveyed *category-diagnostic information* (filled rectangles, with color indicating the chirp classes shown in **A**) defined by underlying acoustic structure of three unique chirps drawn from the same chirp class (consistent color). The other, simultaneous, band (open dotted-line rectangles) had no underlying structure because the three chirps were drawn from 3 (of the 4 possible) distinct chirp classes (different colors). For Categories A and B, the high frequency band was diagnostic; for Categories C and D the low frequency band was diagnostic. The insets on the right show a detailed example from Category B (high-band-diagnostic, Chirp 2 structure) and Category D (low-band-diagnostic, Chirp 4 structure) as time x frequency spectrograms with color overlays indicating chirp class for visualization. **(C) Categorization Training.** Across Days 1-5, an *Alien Identity+Size* participant group (N=95, with N=49 reaching criterion expert performance) learned to associate diverse exemplars drawn from the alien identity categories with one of four distinct “space aliens” (see **B**, left). Two-alternative forced choice (2AFC) trials (blocked by high-versus low-frequency diagnostic band, with feedback) were interleaved with 4AFC trials with novel exemplars (no feedback) to assess generalization of learning. On Day 1 and Day 5, these participants learned to categorize exemplars drawn from all four alien identity categories as “big” or “small” aliens, according to exemplar amplitude (independent of alien identity). In a single session, a separate *Alien Size* group (N=20) trained only on this size judgment without training on alien identity.

Within these maps, functional MRI studies have shown that stable, stimulus-driven voxelwise frequency selectivity can be established across much of auditory cortex (Moerel, de Martino, & Formisano, 2014). Voxels’ relative frequency preference can also be mapped by *explicit attention to specific frequency bands* within a complex, dynamic, and full-spectrum acoustic scene (da Costa et al. 2013; de Martino et al., 2015; Dick et al., 2017; Riecke et al., 2017). The degree to which activity in cortex can be similarly driven by attention to a particular frequency band and ‘bottom up’ acoustic stimulation in that same frequency band can be assessed by constructing ‘concordance maps’ (Dick et al., 2017). These maps quantify the similarity of stimulus- and attention-driven tuning of the subset of voxels within each of a number of small parcellation units within auditory cortex. Such concordance maps allow one to make group-level spatial inferences about where there are consistent relationships between stimulus- and attention-driven frequency tuning in auditory cortex, while taking into account the known cross-participant variability in the exact position, size, and orientation of tonotopic maps relative to gyral morphology (Moerel, de Martino, & Formisano, 2014). We use this analytic approach to ask whether, like overt attention, the presence of category-diagnostic information in voxels’ preferred frequency band drives relatively greater activation, compared to when category-diagnostic information is conveyed via the less-preferred frequency band. Such cross-task similarity would support hypotheses that with learning and expertise, attention is directed toward category-informative dimensions.

As noted above, many naturally occurring sounds convey information about multiple types or categories of information, and this information is conveyed along different acoustic dimensions. To model this, we created all sound exemplars not only to convey ‘alien identity’ information within one of two frequency bands, but also to simultaneously convey ‘alien size’ information via perceived loudness. Here, lower amplitude sounds are diagnostic of ‘small aliens’ and higher amplitude sounds are diagnostic of ‘big aliens’. Our main group of participants trained over five days to learn alien identity (spectrotemporal dimension) and trained, as well, to categorize the same sound inventory according to alien size (amplitude dimension). Another smaller group trained *only* on alien size (amplitude) thereby not gaining expertise in classifying alien identity according to the spectrotemporal information.

This approach allows us to answer several central, yet open, questions. Is category-relevant perceptual information prioritized relative to simultaneous uninformative dimensions? If so, does this selectivity persist when the diagnostic dimension is irrelevant to the task at hand? Do emergent patterns of prioritization align closely with neural activity associated with explicit, sustained attention to the dimension as conveyed by different sounds? Finally, how do these patterns of neural activity relate to the distinct trajectories that learners take in reaching expert-level categorization?

## Results

### Category expertise develops across training, with generalization to novel exemplars

A group of young adults (N=95, see **Methods**) trained for 5 days to develop expertise in rapidly and accurately categorizing complex, multidimensional sounds they had never previously encountered. In the main ‘alien identity’ training task, four sound categories were associated with the identity of four distinct ‘space alien’ images (**Fig 1B**). Each sound possessed acoustic energy in two non-overlapping acoustic frequency bands (∼80-750 Hz and ∼1000-9000 Hz). Both bands were populated with three 400-ms ‘chirps’ played one after another, each varying in frequency contour (**Fig 1A**). In the *category-diagnostic* band, these acoustically variable chirps possessed an underlying regularity that defined category membership (alien identity). Simultaneous and temporally aligned chirps in the other, *non-diagnostic* frequency band were also acoustically variable but possessed no coherent regularity. Category learning thus depended on discovering acoustic regularities shared by category-consistent chirps within a specific frequency band and associating those band-specific regularities with an alien identity. Two categories carried diagnostic information in the high frequency band; two carried information in the low frequency band (**Fig 1B**). This novel soundscape allowed us to capitalize on auditory cortical frequency selectivity to establish cortical regions potentially relevant to the frequency delimited category-diagnostic patterns signaling category identity.

Alien identity categorization training involved a two-alternative forced-choice (2AFC) task, with trials blocked by categories with high- vs. low-frequency category-diagnostic information. Trial-by-trial feedback indicated the correct alien identity category (**Fig 1C**, see **Methods**). After each training block, a four-alternative forced-choice (4AFC) task with no feedback assessed generalization of category learning to novel exemplars not experienced in training (**Fig 1C**). A separate training task involved categorizing exemplars drawn from all four alien identity categories as “big” or “small” according to stimulus amplitude (which varied along a continuum), *without* regard to the spectrotemporal dimensions associated with alien identity (**Fig 1C**, see **Methods**). Two independent samples of participants completed training, one learning alien identity and size categories (*Alien Identity+Size*) and the other learning only size categories (*Alien Size*). This permitted us to test whether identity-training-induced changes in cortical activation associated with the frequency band carrying identity-diagnostic information exist, whether these changes are lacking in identity-naïve listeners, and whether they persist when task demands are redirected to category decisions based on amplitude The *Alien Size* group is also important for identifying any bottom-up salience differences that may differentiate sounds from specific alien identity categories, independent of category expertise.

A subset of expert *Alien Identity+Size* participants (N=49) who achieved at least 75% 2AFC accuracy in reporting alien identity across both high- and low-frequency diagnostic bands by Day 5 returned an average of 9 days later for a single functional magnetic resonance imaging (fMRI) session. Overall, these experts exhibited above-chance 2AFC alien identity training performance on Day 1, t(48)=15.7, p = 7.35 x 10^-21^, with improvements from Day 1 to Day 5, t(48) = 7.82, p = 2.03 x 10^-10^, to reach near-ceiling alien identity categorization (95.5%; SE=0.7%, **Fig 2A, left panel**). Among these experts, 4AFC generalization to novel exemplars exceeded chance on Day 1, t(48)=5.64, p = 4.41 x 10^-7^; M=68.5%, SE=3.3%) and improved significantly from Day 1 to Day 5 (M=87.7%, SE=1.9%, t(48) = 6.56, p = 1.73 x 10^-8^; **Fig 2a, middle panel**). Confusion matrices (**Fig 2B)** from the 4AFC generalization task on the first and last day of training show that category identification sharpens over training. On Day 1, expert listeners made errors in identifying novel exemplars of all categories, and tended to confuse novel exemplars that shared a diagnostic band. By Day 5, listeners’ categorization of novel alien exemplars had sharpened considerably across the board. In addition, the learning trajectories across which alien identity experts reached criterion expertise were heterogeneous; we used this variability to subgroup learners for further analysis (see below).

**Figure 2.**
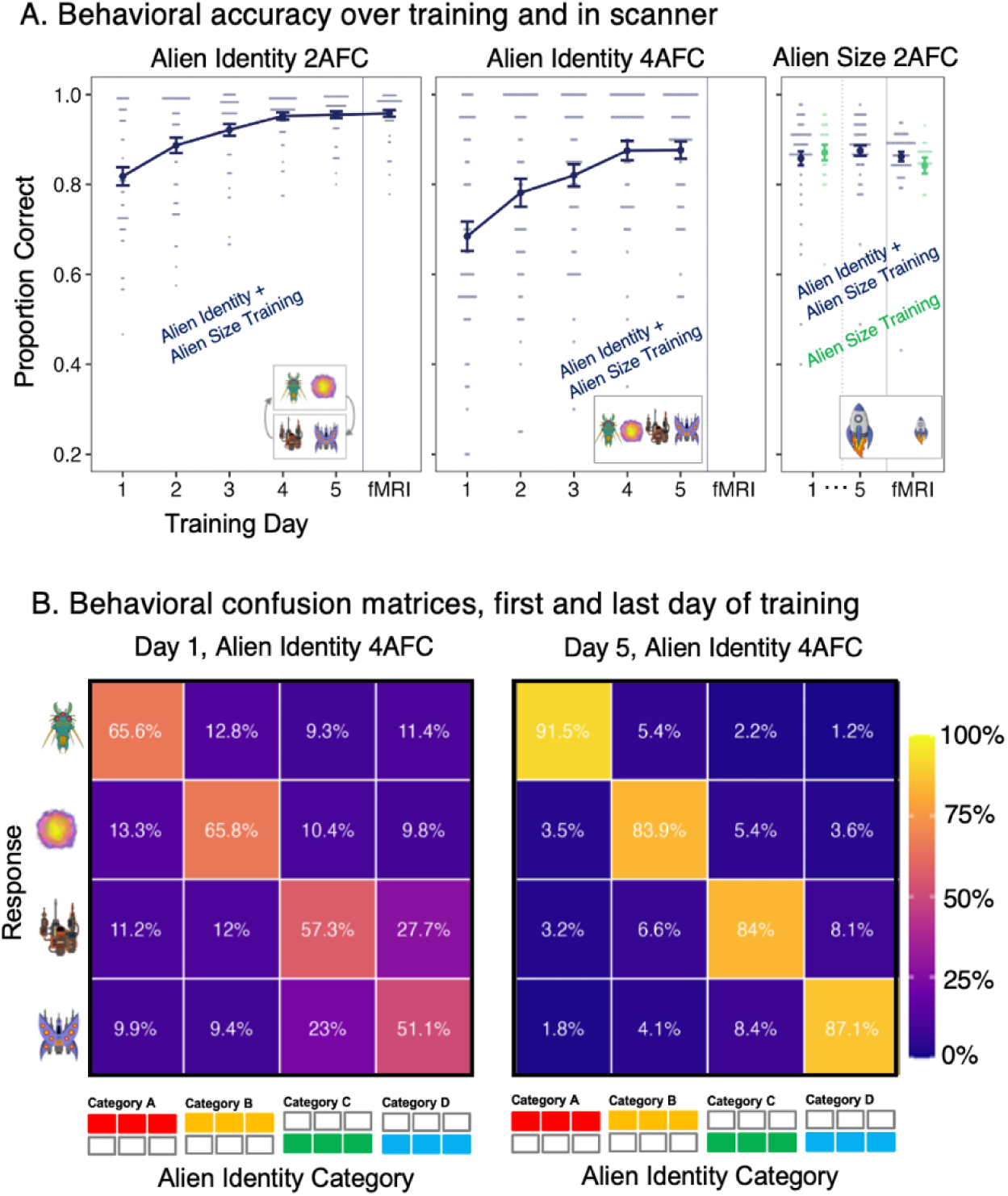
Behavioral Data. **(A) Alien Identity Training (2AFC with feedback).** Proportion correct 2AFC categorization across Days 1-5 (and in the fMRI scanner, no feedback) for the N=49 *Alien Identity+Size* participants that achieved 75% accuracy across all alien identity categories by Day 5. **Alien Identity Generalization (4AFC no feedback).** The same *Alien Identity+Size* participants’ 4AFC categorization of novel category exemplars improved across Days 1-5. **Alien Size Training (2AFC with feedback).** A separate training task involved categorizing exemplars drawn from all 4 alien identity categories as “big” or “small” according to stimulus amplitude. The *Alien Identity+Size* participants who trained on alien identity trained on alien size on Day 1 and Day 5 of training, and in the scanner without feedback. A separate *Alien Size* group (N=20) never trained on alien identity; they trained on alien size for one day and performed the same task in the scanner with no feedback. **(B) Confusions in 4AFC Alien Identity Generalization.** Confusion matrices comparing category ground truth with response for Day 1 versus Day 5 reveal increasing category generalization and specificity. Like Fig 1, x-axis rectangles for confusion matrices show diagnostic frequency band for each category (warm colors: high band, cool colors: low band).

Alien identity experts also trained on the alien size task on Days 1 and 5 to a similar level of accuracy (M=87.5%, SE=1.2%; **Fig 2A, right panel**). The *Alien Size* participants (N=20, sample size matched to the highest-performing ‘Early Expert’ subgroup of *Alien Identity+Size* group, see below) achieved 87.1% (SE=1.7%) alien size categorization accuracy at the end of one training session (**Fig 2A, right panel**).

Both groups took part in a single fMRI session following training. In-scanner behavior echoed patterns of training performance (**Fig 2A, right panel)**. Those who trained on both alien identity and alien size successfully categorized alien identity according to patterns evolving in the diagnostic frequency bands (2AFC task, M=95.8%, SE=0.7%) and according to stimulus amplitude in the size task (M=86.2%, SE=1.1%). Participants trained only on alien size performed with similar accuracy (on alien size) during scanning (M = 84.2%, SE = 1.8%), compared to the group trained on both tasks.

### Alien identity learning drives frequency-band-selective activation differences in tonotopic cortex

We first present functional MRI data from the *Alien Identity+Size* participant group trained on both alien identity and alien size tasks. To identify regions likely to differentially respond to category-diagnostic frequency bands, and to test differences in neural responses as a function of categorization task demands, we first characterized each participant’s unique tonotopic organization on a voxel-wise basis across bilateral auditory-responsive cortex. Here, participants listened to concatenated series of 4-tone mini-sequences that stepped periodically over a 60-semitone range (175 to 5286 Hz, two runs ascending and two descending, see Dick et al. 2017 and **Methods**). To maintain task focus, participants performed a 1-back task, reporting infrequent mini-sequence repeats (in-scanner d’: M=3.55, SE=0.08). From these data we compute voxel-wise activation to mini-sequences grouped into the low and high frequency bands that conveyed alien identity information. The difference in beta coefficients across high-minus-low frequency bands quantifies voxels’ frequency selectivity. As expected from prior work (Moerel, de Martino, & Formisano, 2014), we observe coarse alignment of tonotopic organization across individuals (**SI Fig S1**), but cross-individual heterogeneity in the mosaic arrangement of fine-grained frequency selectivity.

The ‘concordance map’ approach (Dick et al., 2017) described in the introduction allows us to capture consistent regional differences in responses across participants while retaining frequency sensitivity at a voxel-wise level, despite tonotopic map inter-subject heterogeneity. Here, each individual’s auditory-responsive cortices are parcellated into a set of small ROIs defined over the cortical surface (**Fig 3A**, see Methods). Each ROI exhibiting frequency selectivity at the group level (as estimated using the group-average tonotopic map) is sampled into the cortical ribbon in the individual’s native volumetric EPI space (**Fig 3B)**. Analogous to the high-minus-low frequency selectivity contrast from the tonotopy runs, voxel-by-voxel beta coefficients relating to activation during alien identity categorization (involving the high- and low-frequency category-diagnostic bands) are computed and a high-minus-low frequency band difference in beta coefficients is calculated for each voxel. These parallel difference scores permit us to fit a regression model across all participants within each ROI (with participant as a random factor, **Fig 3C**) revealing the ROI-wise concordance between the voxelwise frequency selectivity evoked by tonotopy stimuli, and the hypothesized, category-specific frequency-band selectivity driven by the alien identity learning task (**Fig 3D** and Methods).

**Figure 3.**
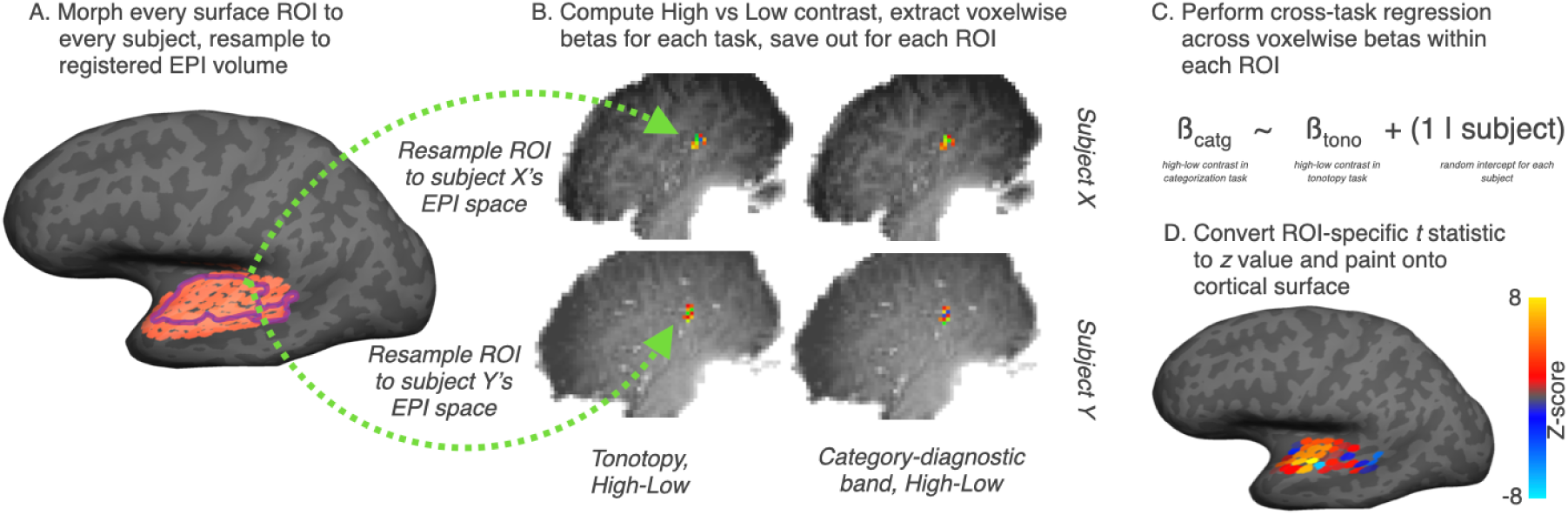
Approach to concordance analyses. To quantify the similarity in activation patterns across tasks, we defined a set of regions of interest (ROIs) and conducted cross-task concordance analyses within each ROI. **(A)** First, we took a set of cortical surface ROIs tiling the superior temporal plane (shown in orange; Dick et al., 2017); for the current work, we only considered ROIs that fell within tonotopically organized auditory cortex as determined by a group average map; these ROIs are outlined in purple. Each ROI was warped to each participant’s native volumetric EPI space. **(B)** For the two categorization tasks, we always examined the contrast between the high-band-diagnostic and the low-band-diagnostic identity categories. We used this contrast even when the task was to categorize stimuli based on overall amplitude, because we hypothesized that, for the *Alien Identity+Size* group, relatively intensive training would result in acquired attentional gain for the identity-diagnostic frequency band, even when performing an unrelated task. For the tonotopy task, we used the contrast between bottom-up sensory input from high vs. low frequencies, and for the attention-o-tonotopy task, we used the contrast between explicit, directed attention attending to high vs. low frequencies in the context of bottom-up sensory input from both frequency bands. Within each volumetric ROI, we extracted voxel-by-voxel beta coefficients for the high versus low contrast in each task. **(C)** To assess the similarity of activation across tasks, we submitted voxelwise betas to a regression analysis for each ROI, separately; the goal in each regression analysis was to predict the beta coefficients from one task using the betas from another. Regressions included by-subject random intercepts. **(D)** The resultant ROI z-statistic, which indicates the strength and direction of the relationship between the two sets of beta coefficients, was painted on an average cortical surface for interpretation and comparison with other maps.

In other words, this approach allows us to ask whether categorization decisions guided by chirp patterns embedded in high versus low frequency bands yield activation in corresponding requency-selective cortical regions. Since each category exemplar possesses rich acoustic energy in each frequency band as well as amplitude variation (signaling size), an exaggerated neural response in the identity-diagnostic versus non-diagnostic frequency band is evidence of expertise-driven changes in cortical activation to the category exemplars.

We first established that these novel ‘alien’ sounds evoked broad activation across much of auditory and auditory-related cortex bilaterally, as well as across a number of other brain networks (**SI Fig S2**). We then turned to the ROI-based regression approach using concordance maps. As seen in **Fig 4A**, we found that, across multiple ROIs, categorization decisions requiring use of information in high versus low frequency bands drives differential activation in voxels exhibiting preferential responses to these acoustic frequencies. Most left hemisphere (LH) auditory cortical ROIs along the swath of the planum temporale – from Heschl’s gyrus posteriorly to the anterior-most aspect – show a strong positive concordance between experience-dependent and category-diagnostic versus stimulus-evoked voxelwise frequency selectivity. Some LH posterolateral ROIs along the superior temporal gyrus (STG) also show positive concordance. In right hemisphere (RH) auditory cortex, we observe significant positive cross-task concordance in multiple clusters of ROIs; in general, FDR-corrected groupings are separated by ROIs showing positive but subthreshold category-tonotopy associations. Across both hemispheres, the few ROIs with negative relationships across tasks tended to be situated in regions exhibiting relatively weak group-level tonotopic selectivity (**SI Fig S1)**. Thus, across expert participants actively engaged in the alien identity category task, we observed that voxels in multiple auditory regions showed greater cortical activation associated with the category-diagnostic versus non-diagnostic frequency band in a way that was predicted by the same voxels’ stimulus-driven preferences for one or the other frequency band.

**Figure 4.**
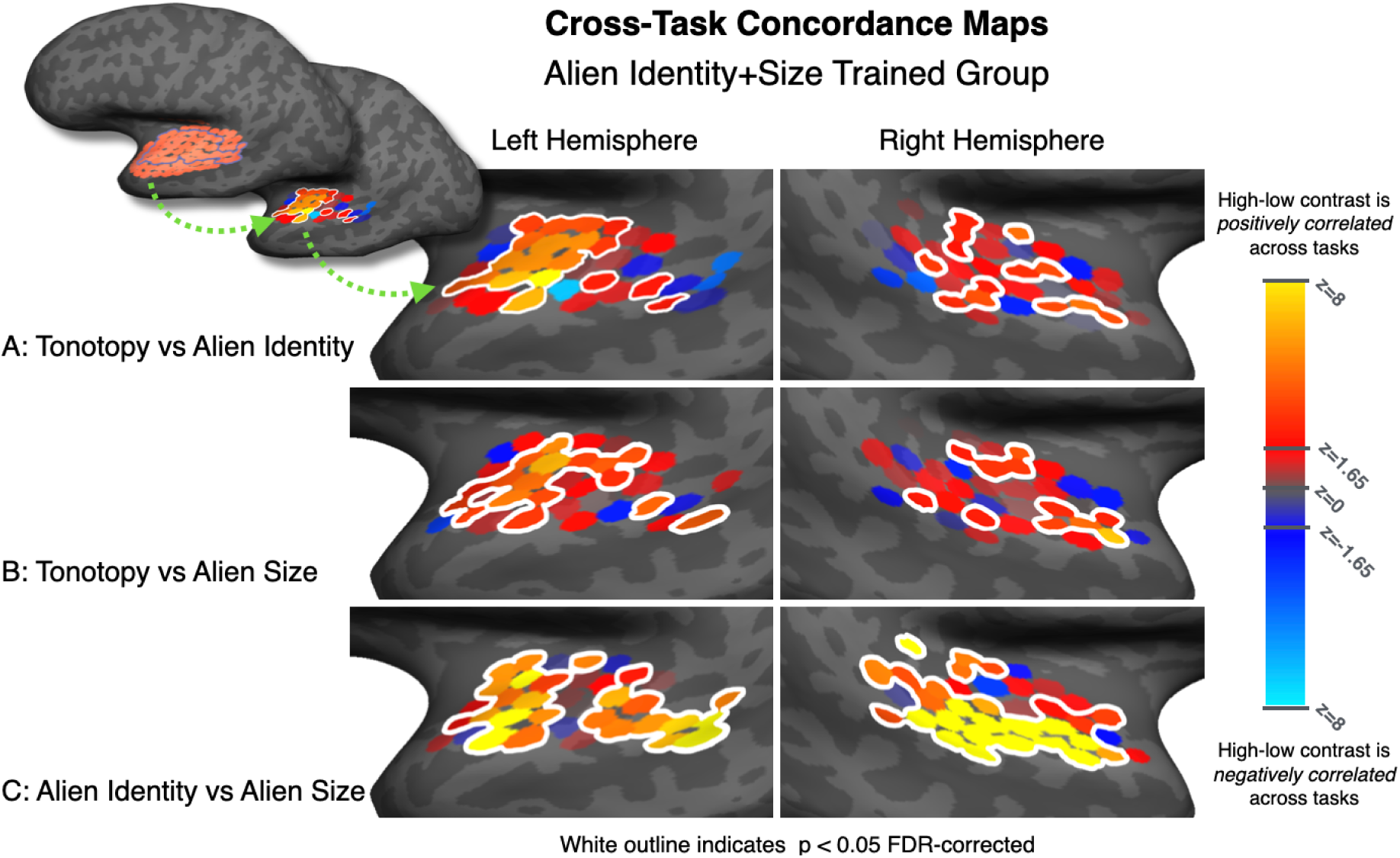
Cross-task concordance maps. Following the approach illustrated in Fig 3, we assessed the concordance among the tonotopy, frequency-diagnostic identity categorization and amplitude-diagnostic size categorization tasks. Each inset figure shows a blow-up of a given concordance map rendered on the left or right temporal lobe. In general, there was a strong positive relationship among betas, indicated by warm-colored ROIs. ROIs outlined in white are statistically significant (p < 0.05 after applying a false discovery rate correction, see **Methods**). **(A)** shows that there is greater cortical activation for category-diagnostic, compared to non-diagnostic, frequency bands during alien identity categorization among the *Alien Identity+Size* group. **(B)** shows that, strikingly, this pattern of activation persists even when categorization according to alien size demands reliance on an orthogonal dimension (amplitude). **(C)** shows that voxelwise selectivity for high-versus low-diagnostic frequency bands also correlates strongly across the identity and size categorization tasks, even though size categorization requires reliance on amplitude.

### Greater cortical activation in voxels preferring category-diagnostic frequency bands persists even in behavioral contexts that do not demand this information

We hypothesized that learning may cause identity-diagnostic dimensions to acquire salience, such that frequency-selective recruitment may persist even when task demands shift to categorization across an orthogonal dimension. To test this, in different blocks, participants also categorized exemplars from all four alien categories according to whether they were “big” or “small” aliens based on amplitude. (Note that sound exemplars varied in amplitude in both alien identity and size tasks, but amplitude was task-relevant only in the size task). As shown in **Fig 2A (right panel)**, in-scanner alien size categorization was accurate, but well below ceiling performance. This indicates that participants successfully relied upon the amplitude dimension for categorization, and that the alien size task was non-trivial.

Although task demands directed participants’ overt attention to an orthogonal acoustic dimension (amplitude), concordance between voxels’ stimulus-evoked frequency preference and exemplars’ frequency-diagnostic band for the over-practiced and more challenging *identity* task persisted (**Fig 4B**). Across all *Alien Identity+Size* participants, in the LH, we observed significant concordance across most of the ROIs in the planum temporale, as well as two ROIs in the posterior STS. In the RH, we observed concordance in three small groups of ROIs in the middle inferior circular sulcus, the anterolateral STG, and two isolated ROIs along the posterior crown of the STG.

Moreover, there was particularly strong concordance between the differences in activation evoked by high- and low-diagnostic band exemplars during *identity* categorization task blocks, and *size* categorization blocks (**Fig 4C**). In the LH, all but the central band of ROIs and a very few of the most peripheral ROIs showed high concordance. In the RH, there was particularly high concordance in ROIs along the entirety of the STG, along with strong concordance in the posterior and anterior inferior circular sulcus.

The differential cortical activation across the high versus low frequency bands cannot easily be attributed to ‘bottom-up’ acoustic salience, as all stimuli possessed information in each frequency band in addition to amplitude variation. Rather, this is consistent with an ‘acquired salience’ for category-diagnostic dimensions that reflects stable changes in cortical activation elicited by category exemplars that persist across changing task demands. Prioritization of category-diagnostic dimensions is present even in contexts that do not demand reliance on them.

### Cortical activation reflects individuals’ trajectories of learning

All *Alien Identity+Size* participants who completed the fMRI scan (N=49) had achieved high levels of expertise in a novel domain. Yet their paths to expertise varied (**Fig 5A**). To understand how these learning trajectory differences might affect neural representations, we grouped participants based on the speed and depth of learning over behavioral training. Early Experts (N=17) achieved at least 90% accuracy on Day 1 for 2AFC training on both high- and low-frequency diagnostic bands. Late Experts (N=16) started at lower accuracies but showed an accuracy increase of at least 20% over the course of 2AFC training and achieved accuracy similar to the Early Experts at the end of training.

**Figure 5.**
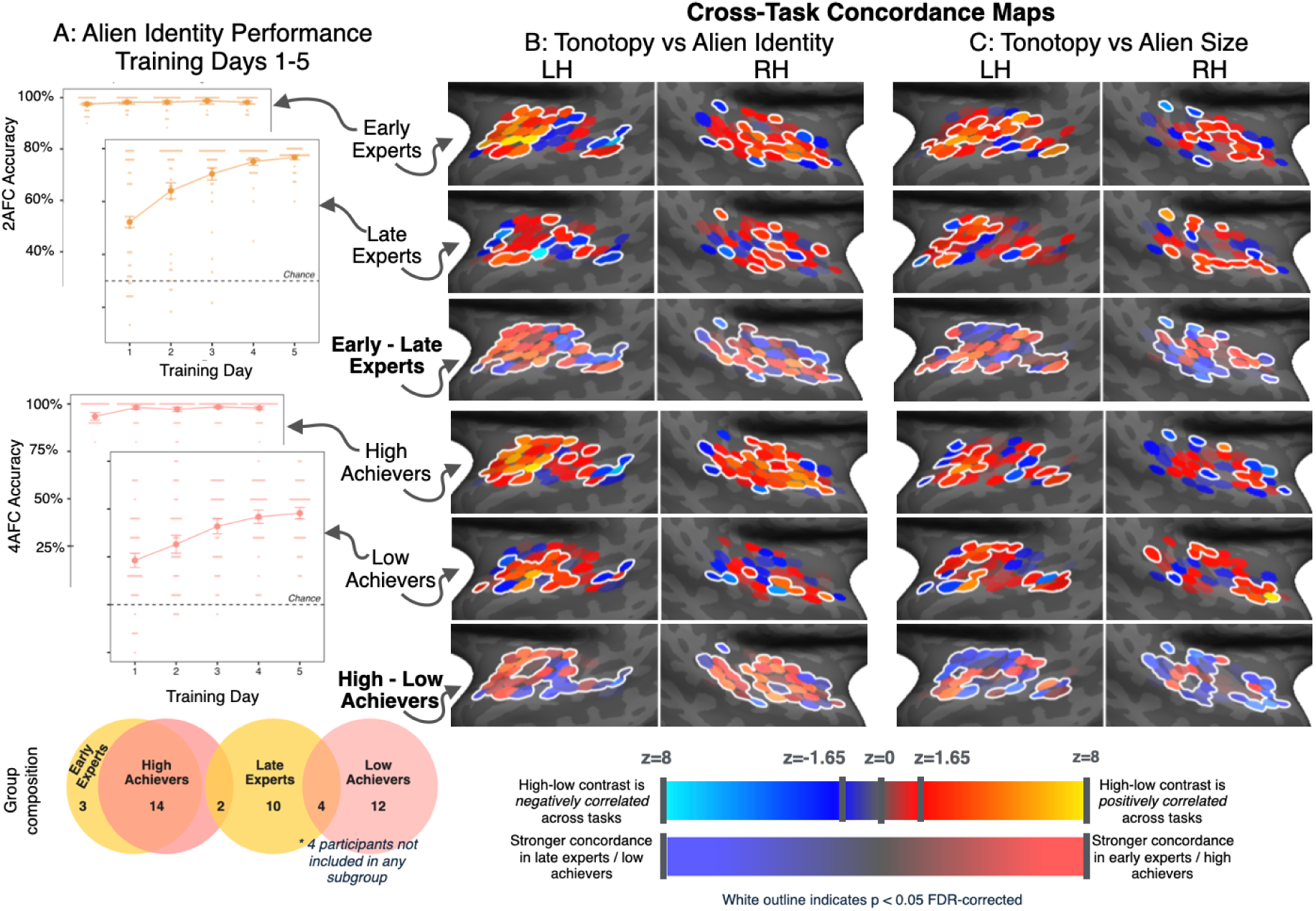
**(A)** Sub-group analyses tested for differences in cross-task concordance as a function of frequency-diagnostic categorization performance during the course of training. One set of analyses considered differences as a function of how quickly participants mastered the 2AFC categorization task: Early experts demonstrated mastery for both high-band-diagnostic and low-band-diagnostic categorization on Day 1, whereas late experts demonstrated substantial improvement over the course of training. A second set of analyses considered differences as a function of generalization ability, separately analyzing data from the top tercile (highest achievers) and bottom tercile (lowest achievers) on the 4AFC generalization task. Behavioral performance and Venn diagrams for these sub-groups are shown in the left panel. Light dots indicate individual participants (no datapoints are obscured by graph overlay), solid points indicate mean performance for each day, and error bars indicate standard error. **(B-C).** We assessed cross-task concordance between the tonotopy task and the alien identity (**B**) and alien size task (**C**); the top two rows show Early and Late Experts concordance maps, with the third row showing the difference between Early versus Late Experts maps. Parallel concordance maps for High and Low Achievers are shown in the bottom three rows. In the difference maps, warm colors indicate ROIs where stronger cross-task concordance was seen in early experts/highest achievers, and cool colors indicate ROIs where stronger cross-task concordance was seen in late experts/lowest achievers.

We also estimated ultimate depth and generalizability of learning by creating separate groups based on generalization performance in the 4AFC task, with the High Achievers (N=16) and Low Achievers (N=16) defined across the top and bottom generalization terciles. These paired subgroups capture (partially overlapping) sets of participants with distinct learning trajectories.

We observe the fingerprints of these trajectories on subsequent cortical responses to category exemplars. Compared to Late Experts, Early Experts exhibit stronger concordance between alien-identity-categorization-driven and stimulus-driven voxelwise frequency selectivity in almost all anterior LH ROIs, particularly the anterior-most row of ROIs located in the inferior circular sulcus (**Fig 5B, top three panels**). In the RH, Early Experts show greater concordance in ROIs along much of the crown of the STG, and somewhat less concordance medially, compared to Late Experts.

Similarly, the High Achievers exhibited greater concordance than the Low Achievers in LH anterior ROIs, as well as somewhat less concordance in the lateral-most posterior ROIs abutting the superior temporal sulcus (STS) (**Fig 5B, bottom three panels**). In the RH, High Achievers showed substantial concordance across almost all ROIs, whereas the Low Achievers – while still expert – showed significant concordance only along some of the lateral STG, and significantly lower concordance than the High Achievers across almost all RH ROIs apart from a few peripheral ROIs.

This indicates that, despite their shared ultimate success in learning and in-scanner task performance, *Alien Identity+Size* participants’ *degree and rate of learning* of identity category expertise is strongly related to the emergence of specific patterns of brain activation that reflect the degree of tuning to category-diagnostic perceptual dimensions.

### Learning-driven attentional gain for a diagnostic dimension depends on the demands of learning

As described above, expert-level categorization drove activation of auditory areas tuned to the frequency band diagnostic of alien identity, even when *Alien Identity+Size* experts were successfully categorizing alien size based on stimulus amplitude. To make a critical test of the importance of learning to weight or attend to the identity-diagnostic frequency band in driving these results, we compared the *Alien Size* trained group (N=20), which remained naïve to alien identity and *never* trained to learn categories based on frequency-band-delimited acoustic information to the Early Expert subset (N=17) of the *Alien Identity+Size* trained group.

As would be expected, alien-size-trained participants exhibited a qualitatively equivalent degree of frequency selectivity and tonotopic organization in auditory cortex (**Fig S1**), relative to the Early Expert group that trained on both alien identity and size. They also showed high behavioral accuracy on the tonotopic mapping task (in-scanner d’: M=2.68, SE=0.13). Having established typical patterns of tonotopic mapping, we next tested whether alien-size-only-trained participants’ auditory cortex would show differential activation for stimuli with identity-disambiguating information in high and low frequency bands when they were performing the amplitude-based alien size task. If there is a systematic activation preference for the alien-identity-diagnostic frequency band among participants trained only on alien size – particularly with a similar pattern of sensitivity to alien identity experts – this would suggest that the spectrotemporal regularities present in the identity-diagnostic frequency band are driving at least some of the results we have observed with alien identity experts, potentially via learning-induced attentional capture (Zhao, Al-Aidroos, & Turk-Browne, 2013).

Though the alien-size-trained participants demonstrated high behavioral accuracy on amplitude-based size categorization during scanning (M: 84.2%, SE: 1.8%, **Fig 2A right panel**) they exhibited little sign of activation for the identity-diagnostic frequency band that matched underlying tonotopy, with only a few ROIs showing significant concordance, and only in the most peripheral regions that are inconsistently and weakly frequency-selective across participants (**Fig 6A**). By comparison, as shown previously in **Fig 5C (top right panel)**, the Early Experts trained on both identity and size tasks showed strong concordance across ROIs between the difference in activation to high-versus low-frequency *identity-diagnostic* band stimuli during alien-size categorization blocks, and high-versus low-frequency tone mapping. A direct statistical comparison between the two groups confirmed that most ROIs showed significantly greater frequency-selective modulation in the amplitude-conveyed alien size task among participants trained on both alien identity and size, compared to those trained on alien size only (**Fig 6B**). This adds further support for the hypothesis that learning categories that are dependent upon statistically structured auditory information restricted to one frequency band drives the acquired salience of that structured information, such that identity categorization experts (but not identity-naïve participants) exhibit frequency-band-specific recruitment of auditory cortex, even when categorizing stimuli along an orthogonal dimension (stimulus amplitude).

**Figure 6.**
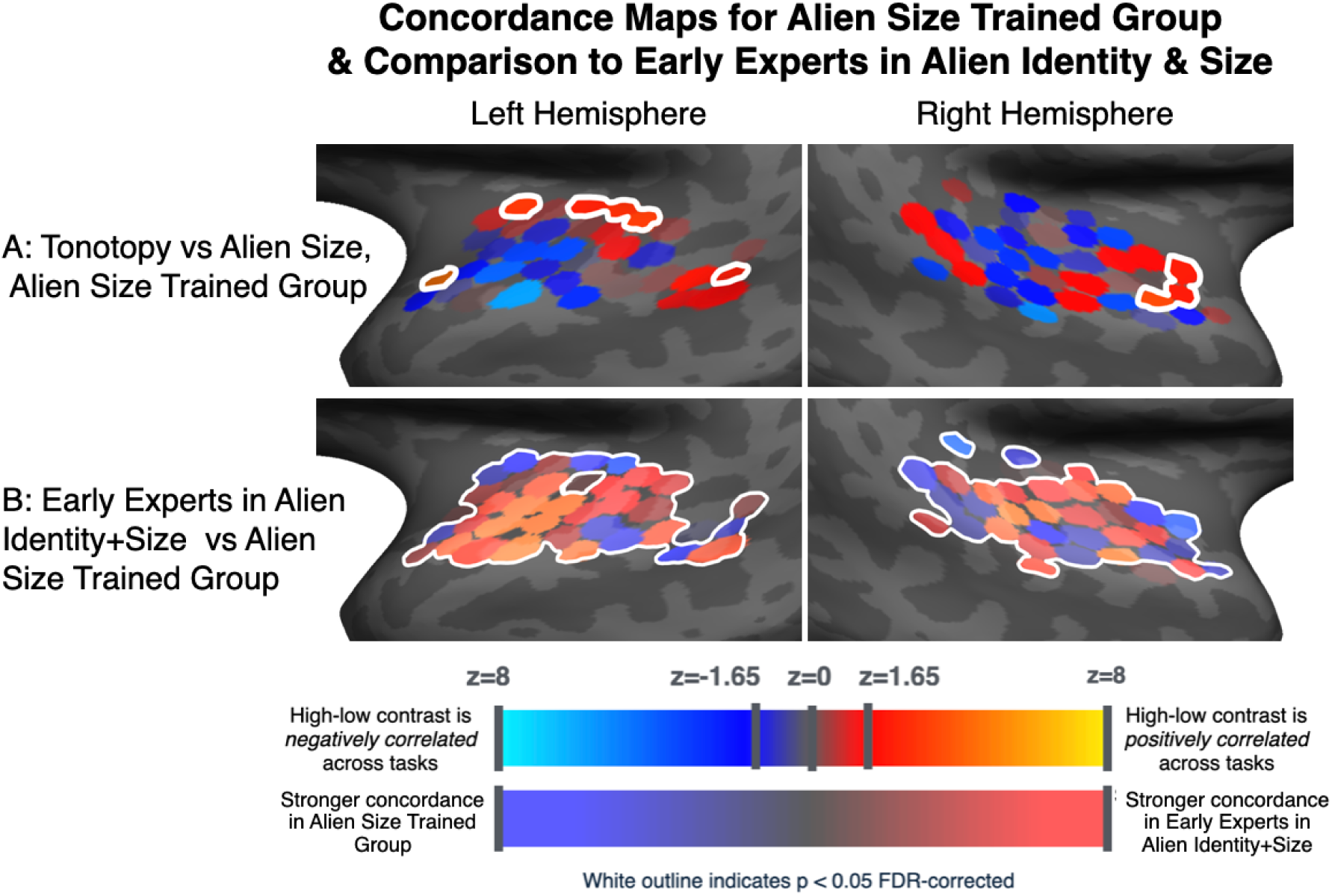
Concordance Maps for Alien Size Trained Group and Comparison to Early Experts in Alien Identity+Size. **(A)** shows the *Alien Size* trained participants (N=20) concordance map between high-versus-low-frequency responses from tonotopy, and response differences in the space alien size fMRI task when comparing activation for alien sounds with *alien identity* information in high versus low spectral bands. (Recall that this participant group was naïve to alien identity.) With the exception of a few medial ROIs, participants did not recruit auditory cortex in a frequency-selective way when performing amplitude-based categorization. **(B)** shows the difference between the analogous concordance map for the Early Expert group of *Alien Identity+Size* trained group **(top right panel of** Fig 5c) versus the *Alien Size* only trained participants. This difference map suggests that the frequency-selective recruitment of auditory cortex that *Alien Identity+Size* experts exhibited in the amplitude categorization task was not driven by properties of the acoustic signal *per se*.

### Frequency-selective activation patterns driven by category learning align with those evoked by overt attention

We have observed that when categorization implicitly hinges on information in a delimited frequency band, voxels more responsive to this band show increased activation compared to when diagnostic information is conveyed via the less preferred frequency band, despite simultaneous acoustic energy in each band. Motivated by theoretical accounts suggesting that that perceptual weighting reflects attentional gain, we next asked whether the pattern of increases in activation for category-diagnostic frequency bands is aligned with activation evoked by *explicit* attention to the same frequency band. Among participants trained on *Alien Identity+Size*, we compared the activation difference between high- and low-frequency identity-diagnostic bands with the parallel activation differences evoked by explicit, sustained attention to one of two simultaneously presented streams of sinewave tones situated within the same high- and low-frequency bands.

In addition to the tonotopy and alien categorization tasks, the *Alien Identity+Size* participants also completed a sustained auditory selective attention task based on the approach of Dick et al. (2017) who found that overt attention directed at one of two sequences of tones positioned in specific frequency bands elicits an attention-driven map across auditory-responsive cortex. As shown in **Fig 7A**, participants heard two simultaneously presented sequences of sinewave tones composed of concatenated 4-tone mini-sequences that varied in amplitude; mini-sequences were played simultaneously in higher and lower frequency bands, corresponding to the diagnostic bands of the alien identity task (see Methods). At the beginning of each task block, explicit instructions directed participants to detect mini-sequence repeats in the high frequency band, the low frequency band, or repeats of the overall relative amplitude of the mini-sequence (1-back task d’: M=2.28, SE=0.13). As anticipated given previous studies of frequency-selective attention (Da Costa et al., 2013; de Martino et al, 2015; Dick et al 2017; Riecke et al., 2017), voxelwise differences in activation when attending to high versus low frequency bands were strongly associated with stimulus-evoked frequency preference (from the tonotopy scans) across most LH auditory cortex ROIs except in the anterior lateral and posterior STG (**Fig 7B**). In the RH, spectrally-directed attention evoked similar activation differences as tonotopic scans along the lateral planum temporale and STG.

**Figure 7.**
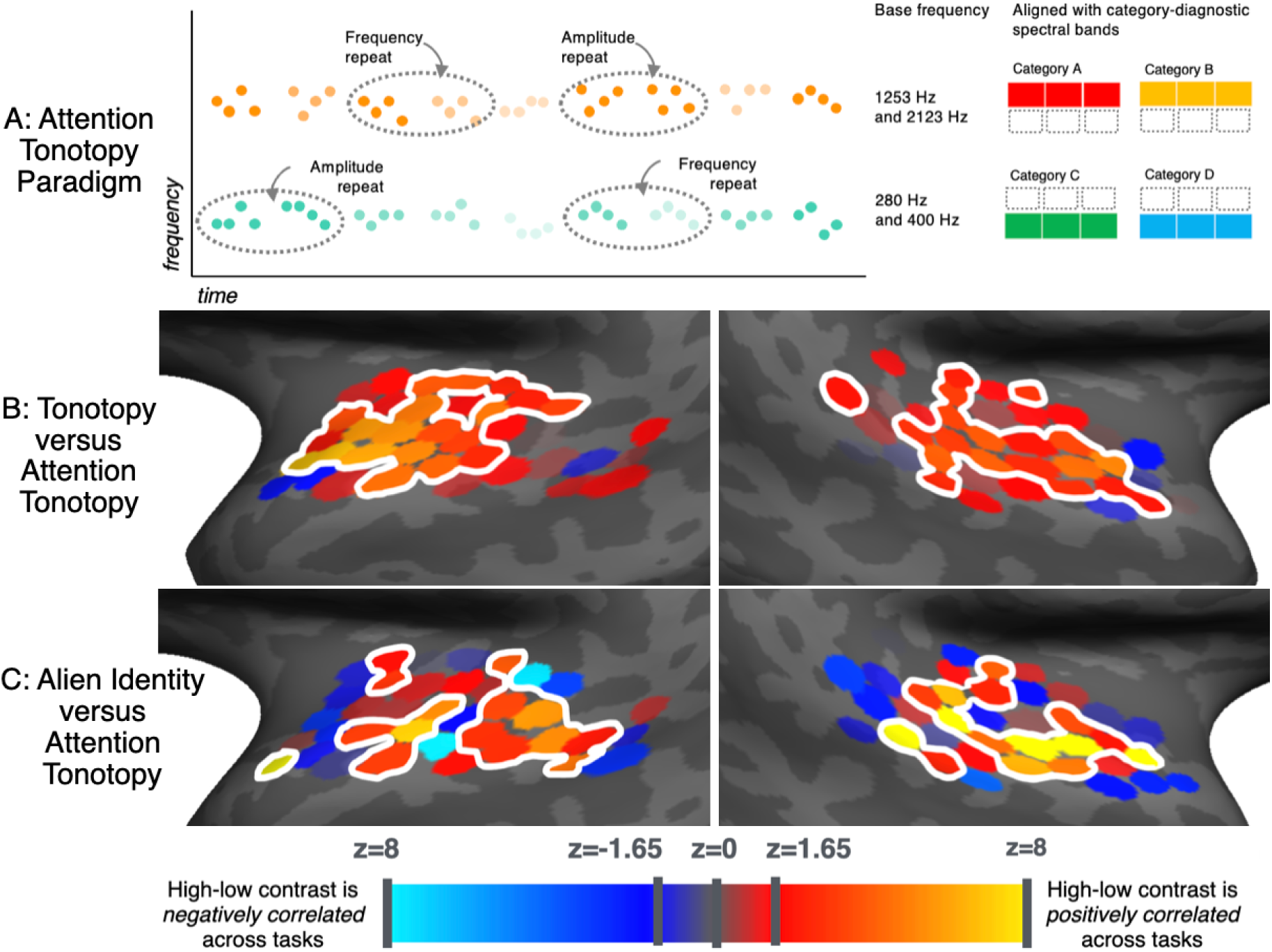
Frequency-selective activation patterns driven by category learning align with those evoked by overt attention. Theoretical accounts suggest that the neural selectivity that emerges with learning is driven by increased attention to diagnostic dimensions. To test this hypothesis, we assessed cross-task concordance with the attention tonotopy task **(A)**, where participants were presented with two streams of short tone sequences, where each stream was in a high- or low-frequency band corresponding to the high and low identity-category-diagnostic frequency bands in the *Alien Identity+Size* task. Participants were instructed to detect repeated 4-tone sequences in one or the other band, depending on block. Tone sequences also differed in overall amplitude (indicated by transparency level). **(B)** Analyses considering all participants showed strong concordance between attention-driven activation to high versus low frequency bands and the high-versus low-frequency band activation difference elicited by the tonotopy task (i.e., stimulus-driven activation). **(C)** Concordance between alien identity categorization-driven activation to high versus low diagnostic frequency bands, and explicit attention to the same frequency bands populated with simple sequences of tones was observed bilaterally along much of the lateral aspect of tonotopically mappable auditory cortex.

We then asked whether patterns of cortical activation elicited by *explicit, directed attention* to the frequency bands align with activation patterns that *arise in category decisions that are implicitly reliant on information* in these same frequency bands. As shown in **Fig 7C**, cross-task concordance maps showed that analogous differences in activation to high versus low frequency bands were observed laterally in both hemispheres, in a swath of ROIs along the central portions of the STG and abutting ROIs in the planum temporale.

In summary, the more lateral aspects of auditory cortex showing differential activation when listeners explicitly attend to a delimited region of the frequency spectrum show correspondingly differential activation when categorization *implicitly* hinges on information situated in these same frequency bands. This holds true despite the considerable acoustic differences between the sequences of pure tone stimuli used for the explicit attention task and the spectrotemporally more complex and structured alien sound category exemplars.

## Discussion

How does perception come to weight the most diagnostic patterns of input for the task at hand? We used fMRI to probe the neural consequences of learning to categorize across a novel soundscape defined by multiple, complex acoustic dimensions evolving in time. To do this, we situated category-diagnostic patterns into frequency-delimited bands thereby coarsely segregating category information into different channels in tonotopically organized auditory cortex. This allowed us to make targeted predictions about: a) how driven modulation of cortical activity would intersect with existing tonotopic functional regionalization; b) how different trajectories of learning to expert-level performance would be reflected in these representations; and c) how a neural signature of ‘learned salience’ and/or attentional gain for the category-diagnostic information might align with overt attention directed toward distinct stimuli in the same frequency bands.

We observe that category learning drives prioritization or gain across a category-relevant dimension. This is reflected in greater cortical activation in voxels whose frequency response is associated with the category-diagnostic frequency band, compared to the simultaneously presented non-diagnostic band. This prioritization persists even when the in-scanner task shifts to categorization based on a completely different auditory dimension, amplitude. Further, it partially aligns with the patterns of cortical activation that emerge when listeners explicitly direct sustained attention to unrelated sounds situated in the same frequency band. This is consistent with ‘attentional gain’ accounts of advantaged perceptual processing of category-diagnostic dimensions. These neuroimaging results support the notion that learning complex categories can drive the emergence of acquired attentional gain implicitly directed to category-relevant perceptual dimensions.

The question of whether - let alone how - sensory cortical representations are impacted by the acquisition of category expertise has not had a clear answer. Evidence of cortical modulation with category expertise comes predominantly from visual cortex under conditions of active categorization of static images (Sigala and Logothetis, 2002; De Baene et al. 2008). However, these results are mixed, with some studies not observing effects of categorization on visual cortex (Freedman et al. 2003; Jiang et al. 2007; Gillebert et al. 2008; van der Linden et al. 2010) potentially consistent with category representations arising in flexible, more ‘amodal’ cortical areas with little lasting influence on sensory cortex (Freedman et al. 2003; Serre et al. 2007; Roy et al. 2010, but cf. Hausfeld, Riecke, & Formisano, 2018). Jiang et al. (2018) report auditory category learning results consistent with this two-stage model, and others have made a case for prefrontal involvement in auditory categorization (Russ et al., 2007). In contrast, our results indicate stable, training-induced changes in auditory cortex that persist even when task demands direct attention away from the category-diagnostic dimension (see also Folstein et al. 2013 and Van Gulick & Gauthier, 2014 for evidence from repetition suppression across visual categories).

These changes were driven by the specific demands of learning, and not by mere exposure to statistically structured sounds. Participants who trained to use frequency-band-diagnostic information to categorize alien identity exhibited neural selectivity consistent with acquired attentional gain to the diagnostic frequency bands. But those who learned the amplitude-signaled size categories across the very same sounds, thus experiencing the identity-diagnostic spectral information without learning to use it to categorize, did not. Although humans and other organisms exhibit an ability to learn across passive exposure to statistically structured input (Saffran & Kirkham, 2018; Wilson et al., 2015), multiple studies have observed that complex acoustic exemplars that model the complexity of speech are not well learned over passive exposure (Wade & Holt, 2005, Emberson, Liu, & Zevin, 2013). Yet alignment of these statistically structured sounds with active behavior – even incidentally and without overt feedback – can rescue category learning (Roark, Lehet, Dick, & Holt, 2022). The cross-group differences in the present study demonstrates that the alignment of feedback with a diagnostic dimension played a role in eliciting the cortical selectivity toward those dimensions. These patterns of acquired attentional gain in human auditory cortex connect with nonhuman animal electrophysiological studies that observe that the task-relevance of an acoustic dimension drives changes in receptive fields consistent with attentional gain (Atiani et al. 2014; Bagur et al., 2018).

In this vein, Yin et al. (2020) present evidence for top-down modulation of auditory cortex by frontal cortex in ferrets, along with induction of intrinsic auditory cortical patterns related to acquired categories that persist even in passive listening. Though we limited our investigation specifically to tonotopically mappable temporal cortex, the regions we observe to be modulated by category learning receive rich top-down feedback inputs, and exhibit modulation by behavioral context that extends beyond basic representation of perceptual dimensions (e.g., Zhong et al. 2019). Such top-down modulation has been well-documented in nonhuman animal electrophysiology (particularly in the visual system, e.g., Freedman et al. 2003; DeGutis and D’Esposito, 2007) and may interact with the persistent changes to auditory cortical representation arising from category learning that we observe.

In the present study, we see that participants - above all the most expert - show selectively greater activation for the category-diagnostic frequency band. Importantly, this effect is widespread across bilateral auditory cortex, particularly in anterior and lateral portions. This degree of auditory cortical modulation by the category-diagnostic dimension - even when task-irrelevant - is striking by comparison with other studies of sensory cortical category sensitivity which tend to show greater category readout in frontal regions (e.g., Jiang et al., 2007, 2018). In situating category-diagnostic dimensions into discrete frequency band, our approach may have afforded greater sensitivity by aligning learning-driven cortical changes with the primary representational axis of auditory cortex.

The question of whether such category-learning-related modulation of neural activity is a consequence of learned attentional gain to diagnostic dimensions also has surprisingly little direct support, despite the centrality of this proposition in theories of visual (Kruschke & Johansen, 1999; Love, Medin, & Gureckis, 2004; McColeman et al., 2014) and speech (Heald & Nusbaum, 2014) categorization. It is clear that explicit attention to stimulus features modulates representations in both visual (Foster & Ling, 2022; Gundlach et al., 2023; Saenz et al. 2002; Serences & Boynton, 2007; Treue & Maunsell, 1999; Yoo et al., 2022) and auditory (DaCosta et al. 2013; Dick et al. 2017; Hausfeld, Riecke, & Formisano, 2018; Petkov et al., 2004; Riecke et al. 2017) cortical response. Here, we asked whether patterns of cortical gain driven by explicit attention are concordant with patterns that emerge implicitly across category-diagnostic dimensions in categorization. Consistent with our prior research (Dick et al., 2017), sustained auditory selective attention across sequences of tones presented simultaneously in two frequency bands results in greater activation across the attended frequency band. Here, in the same listeners, these patterns aligned with the prioritization of frequency-selective cortical activity driven by categorization demands, but primarily in more lateral auditory cortical regions. Of note, this concordance is not driven by task demands; prioritization of the diagnostic frequency band persists even when task demands change and frequency band is no longer task-relevant. This suggests that prioritization of diagnostic information has become a stable, lasting part of how category exemplars are represented in experts which is absent in category-naïve listeners. To our knowledge, this presents the first empirical evidence linking acquired gain to category-diagnostic dimensions of acoustic input and overt, directed selective attention.

The current results also shed light on what processes might be leading to expertise-driven changes in neural recruitment (Haist, Lee, & Stiles, 2010; McGugin, Gatenby, Gore, & Gauthier, 2012). Often, experts show selectively increased sensory cortical activation for the object of expertise (e.g., Bilalić, 2016; Gomez, Barnett, & Grill-Spector, 2019), but the causal change in neural or cognitive processing leading to this activation can be challenging to ascertain (McKone, Kanwisher, & Duchaine, 2007; Op der Beeck, Pillet, & Richie, 2019). Here we show that selective activation changes in experts are systematically related to the locus of category-disambiguating information, even when that information is not immediately task-relevant. This, along with the relative similarity of category- and attention-selective frequency modulation maps (**Fig 7**), lends support to accounts suggesting learned expertise drives a lasting attentional bias for category-relevant dimensions (Folstein, Palmeri, & Gauthier, 2012). (Alternative accounts emphasize dimensional stretching (Folstein, Palmeri, van Gulick, & Gauthier, 2015, or attention-gated plasticity (Sasaki, Nanez, & Watanabe, 2010).) It is important to note that extensive experience with perceptual categories or objects, such as one’s own name or ringtone, can promote automatized overall attentional capture (e.g., Roye, Schröger, Jacobsen, & Gruber, 2010). However, the neural signature of such global ‘attention-grabbing’ effects will differ from those evoked by dimension-selective attention when the category-diagnostic perceptual dimension is topographically mapped in cortex (for instance across retinotopic space, or acoustic frequency as in the present study), and the presence of category-informative information may be differentially distributed within subregions of this space (Gomez, Barnett, & Grill-Spector, 2019; Norman-Haignere, Kanwisher, & McDermott, 2015).

Disambiguation of different accounts could be furthered through explicit modelling of neural response functions that would efficiently partition category exemplars (e.g., Liu, Montes-Lourido, Wang, & Sadogopan, 2019). Such modelling may help to shed light on why there is greater category-selective modulation of activation in anterior, versus posterior, auditory cortical regions, particularly in the most expert listeners. Additionally, data that disambiguate signals from upper versus lower cortical layers would potentially reveal more fine-grained information about neural encoding of category-relevant information (Montes-Lourido, Kar, David, and Srivatsun (2021) as well the role of learned attention in such encoding (Moerel, Yacoub, Gulban, Lage-Castellanos, & de Martino, 2021; Heynckes, Lage-Castellanos, De Weerd, Formisano,& De Martino, 2023).

In future work, it will be important to show that the increased activation to identity-category-relevant frequency regions that is evident when they are not task-relevant (e.g. during size judgments) is not resulting from carry-over across experimental blocks. While we believe this is unlikely given the difficulty of the amplitude-conveyed alien size task (where weighting by spectrotemporal cues is counterproductive), it is possible that the activation patterns indicating acquired gain of the identity-diagnostic frequency band may diminish when not regularly reinforced via interleaving with blocks of the identity task. Use of a more naturalistic or captivating training paradigm (Deveau, Lovcik, & Seitz, 2014; Lim, Fiez & Holt, 2019; Whitton, Hancock, Shannon, & Polley, 2017) would promote extended training and consolidation, which would be particularly revealing in tracking slower learners like those who did not achieve full mastery in the present study. We also predict increasing learned salience such that identity category experts would exhibit further increases in gain for the identity-diagnostic band in the size categorization task, despite the task-irrelevant nature of frequency-band-delimited alien identity information in this task.

## Methods

### Participants

Our study involved two separate groups. The *Alien Identity+Size* group trained to categorize sounds according to alien identity (based on complex spectrotemporal patterns) and, separately, to categorize the same exemplars according to alien size (based on amplitude). An *Alien Size* group trained only on alien size. All participants were fluent English speakers with normal hearing, had normal or corrected-to-normal vision and no history of neurological impairment or speech/language disorder, and reported no experience with a tonal language.

The *Alien Identity+Size* group included participants from the Pittsburgh, PA, USA and London, UK communities who completed a five-day training protocol (N=95; 18-40 yrs). Individuals who met criterion levels of expertise in alien identity categorization (75% across both diagnostic frequency bands on the 2AFC alien identity task on Day 5) were invited to participate in a subsequent MRI session, with data collected from N=54 experts. Five participants’ data were excluded (motion, non-compliance with task) resulting a final sample of N=49 experts (18-37 yrs, M=23.9 yrs; 29 female, 18 male, 2 non-binary; 59% White; 32 Pittsburgh, 17 London).

The *Alien Size* group (N=20, 18-43 yrs, M=28.1 yrs; 12 female, 8 male; 55% White; 20 London) trained only on the alien size task and participated in an abbreviated MRI session.

### Stimuli

Four auditory categories were sampled from a highly complex perceptual space (see Obasih et al., 2023). As shown in **Fig 1**, each stimulus was composed of patterned acoustic energy in two non-overlapping acoustic frequency bands (∼80-750 Hz and ∼1000-9000 Hz). Each band was populated with three 400-ms tonal ‘chirps’ (100-ms ISI) varying in frequency contour. In the *alien-identity-diagnostic* frequency band (high-frequency for Categories A, B and low-frequency for Categories C, D), the three acoustically variable chirps possessed an underlying regularity that defined alien identity category membership. The three temporally aligned chirps in the non-diagnostic frequency band (low for Categories A, B; high for Categories C, D) were also acoustically variable but possessed no coherent underlying regularity.

To create a large set of naturally varying category exemplars, we derived chirps from four native Mandarin Chinese talkers (two female) uttering single-syllable utterances in each of Mandarin’s four lexical tones, which convey variable, but structured, fundamental frequency (F0) contours. We extracted the F0 contour from each and resynthesized the contour to create a non-speech chirp, time-normalizing each to be 400 ms. We pitch-shifted each chirp +33 semitones, then applied a 1000 Hz high-pass filter to create a pool of chirps to populate the high frequency band. Equivalently, to create chirps for the low frequency band, we pitch-shifted the F0 contour −1 semitone and applied a 500-Hz low-pass filter.

Acoustic information in the category-diagnostic band consisted of three distinct chirps derived from the same talker and the same Mandarin lexical tone category, thereby conveying an acoustically variable but structured regularity in frequency contour. Acoustic information in the non-diagnostic band consisted of three different chirps drawn from three *different* Mandarin lexical tone categories (from among any of the four) such that there was no underlying acoustic regularity across chirps. In all, this created a stimulus pool of 36,000 novel category exemplars (Obasih et al., 2023) from which we sampled 592 unique stimuli.

In addition, we also imposed orthogonal amplitude variation (between 62-80 dB) on copies of each of these 592 sounds. This met two goals: to increase variability and task difficulty, and to permit creation of two *Alien-Size* categories, with "Big Alien" exemplars falling between 72-80 dB sound intensity, and “Small Alien” exemplars falling between 62-70 dB (all reported dB referenced to 2*10^-5^ and measured in Praat). Thus, each exemplar could be categorized along identity (spectrotemporal, frequency-band-specific) or size (amplitude) dimensions.

### Procedure

Online categorization training was implemented using Gorilla software (Anwyl-Irvine, Massonnié, Flitton, Kirkham, & Evershed, 2020), the Google Chrome browser, wired headphones (with compliance screening as in Milne et al., 2021), and a computer/laptop (not tablet/phone). Stimuli were created with Matlab version 2021b (The MathWorks), Praat (Boersma, 2001) and SoX version 14.4.2 (www.sourceforge.net).

#### Behavioral Training Prior to fMRI

*Alien Identity+Size* participants learned to associate 1400 ms novel sounds with one of four alien images via category-informative feedback identifying the correct alien after each response. Each of 5 daily sessions involved two interleaved tasks. The first was a 2AFC alien identity categorization training blocks in which participants indicated identity with a keypress, and feedback (1500 ms) indicated the correct alien. In each 120-trial training block, trials were segregated according to the category-diagnostic high or low frequency band (Categories A vs. B and Categories C vs. D). Every 20 trials, the category pair, and thus diagnostic frequency band, alternated; the other two alien-identity response options were blanked out as response alternatives. Training stimuli were randomly selected from a pool of 512 exemplars (128/category). Amplitude was balanced across trials and roughly equated across categories on each training session. Each training block was followed by a 20-trial 4AFC generalization block with all aliens as response options, and no feedback. These trials involved a separate pool of 20 exemplars/category that were never presented in training to assess generalization of learning (see **Fig 1c**). Across four 2AFC and 4AFC blocks, category exemplars were mixed with a fixed level of scanner noise (57 dB) to familiarize participants with categorization in an MRI environment. The visual position of aliens (and associated keypress) was fixed the first two days of training, then allowed to vary starting on Day 3. (Note that Obasih et al. 2023 found that alien category learning speed and attainment was surprisingly unaffected by different training regimes, including manipulations of exemplar variability, blocking versus interleaving categories in training, and overt instructions to attend to the diagnostic band.)

On Day 1 and Day 5, these participants also trained on alien size categorization. Across the first eight trials, one exemplar from each of the four categories was presented at the highest amplitude (80 dB), then one exemplar from each alien category was presented at the lowest amplitude (62 dB). The amplitude difference diminished each 4 trials for a total of 40 trials, with 71 dB defining the category boundary between “big” and “small” aliens. In a second 40-trial block, amplitude values were fully randomized with equiprobable big/small exemplars and an equal number of exemplars from each alien category. A fixed level of scanner noise (again 57 dB) was mixed with stimuli to mimic the MRI sound environment. Additionally, a separate *Alien Size* participant group completed a single session of 2AFC alien size category training over 80 trials.

#### Tonotopy

To characterize voxel-wise frequency selectivity efficiently for each participant. participants were scanned as they listened to sequences of 178-ms sinewave tones (0 ms ISI, 5 ms ramp at on/offset), organized in 4-tone mini-sequences, each separated by 356 ms. Participants reported infrequent mini-sequence repeats in a 1-back task. The frequency range of the four tones composing the mini-sequences increased or decreased in frequency incrementally in 10 logarithmically scaled steps to sweep across 175-5286 Hz (60 semitones). Across the 60 mini-sequences contained in each 64-sec sweep, 3 mini-sequence repeats occurred quasi-randomly. The swept frequency permitted fMRI analysis using a) phase-encoded mapping, where voxels responsive to a particular frequency should respond at a consistent phase delay (Dick et al., 2012; 2017; Sereno et al., 1995) and b) multiple regression analysis to identify voxels responsive to the range of the lower (< 500 Hz, lowest three steps) and higher category-diagnostic frequency band (≥ 1000 Hz, highest five steps) identity-category-diagnostic frequency bands.

For analysis of training and in-scanner behavior, the response window for each mini-sequence was defined from the onset of the final tone to the onset of the final tone in the subsequent sequence (1068-ms response window). For calculation of d-prime scores, extreme hit and false alarm rates of 0 or 1 were adjusted following the approach of Stanislaw and Todorov (1999). As an introduction to the task, participants familiarized with one up-sweep and one down-sweep on Day 4 of online training (or at the end of the single training session for the *Alien Size* group), with mini-sequence repeats indicated on-screen. As with the space alien task, during out-of-scanner training, scanner EPI noise (57 dB) was mixed in with tone stimuli (∼80 dB).

#### Attention tonotopy

Participants who trained on the *Alien Identity+Size* categories completed a task to characterize voxelwise frequency-selective and amplitude-selective overt attentional responses in individual participants. Following the approach of Dick et al. (2017), participants heard 4-tone mini-sequences evolving simultaneously in two frequency bands with instructions to “attend low” or “attend high” and report mini-sequence repeats from the attended frequency band. To parallel the amplitude differences signaling alien size categories, mini-sequences alternated between higher and lower amplitude levels, which differed on average by ∼10dB. A different spoken instruction directed participants to “attend volume,” reporting 1-back matches on the attended amplitude dimension; data from the "attend volume" condition were not used for the present study. Mini-sequences were composed of 180-ms sinewave tones drawn from a pool of four frequencies (1, 3, 5 and 7 semitones above a ‘base’ frequency). Tones were drawn with replacement, with the constraint that a sequence could never be a single tone played four times. Simultaneous tones from high and low frequency bands were temporally aligned. Mini-sequences were separated by 360 ms silence in short blocks that concluded with 2 sec of silence. Each block included two mini-sequence repeats in the high-frequency band, two repeats in low-frequency band, and two amplitude repeats. The 1080-ms response window for each mini-sequence was defined from the onset of the final tone to the onset of the final tone in the subsequent sequence.

Participants familiarized with the task on Day 3 of categorization training and first experienced mini-sequences in single frequency bands of fixed amplitude. Next, amplitude variability was introduced (first only with the task of reporting tone sequence repeats), then finally, participants practiced with dual-frequency-band amplitude-variable stimuli. On-screen messages alerted participants to mini-sequence repeats in the attended band. There was no scanner noise mixed with stimuli during familiarization. Participants then completed 24 attention tonotopy trials (3 blocks/trial, 11-17 mini-sequences/block), with on-screen feedback provided for the first half of the trials. For half of the trials, the dual-band stimuli had base frequencies of 280 and 1253 Hz, and for the other half, the base frequencies were 400 and 2123 Hz. “Attend high,” “attend low” and “attend volume” trials were equiprobable. The amplitude of the stimuli was set so that the two frequency bands were approximately equated on perceptual loudness (∼73 dB average), and a fixed level of scanner noise (57 dB) was presented throughout these trials.

#### MRI Procedure

Images were acquired in a single 90-min session with 3T Siemens Prisma scanners and 32-channel head coils (CMU-Pitt BRIDGE Center in Pittsburgh, RRID:SCR_023356, and at the Birkbeck-UCL Centre for Neuroimaging in London). Identical pulse sequences were used across sites, and very similar QA procedures were in place by the respective MRI center teams to monitor scanner performance. Head movement among London participants was minimized with a head stabilization prototype (MR Minimal Motion System). Both sites delivered audio over Sensimetrics earbuds with pre-filtering to accommodate the earbuds’ response profile, presented tasks using PsychoPy (v. 2022.2.4), and recorded responses with a button box.

Participants in the *Alien Identity+Size* group who qualified for the MRI session completed a 15-min online refresher one day prior to the scan. This involved two tonotopy trials (one up-sweep, one down-sweep), 80 2AFC categorization trials (48 alien identity, 32 alien size, drawn from the pool of 80 generalization exemplars with no feedback), and two short 9-block attention-o-tonotopy trials with parameters mirroring the MRI task. Participants in the *Alien Size* group completed a refresher within one day of their scan. It included two tonotopy trials and 32 trials of 2AFC alien size categorization, with no feedback.

At the scanning session, structural images were acquired using a T1-weighted magnetization-prepared rapid acquisition gradient echo (MPRAGE) sequence (TR = 2300 ms, TE = 2.98 ms, FOV = 256 mm, flip angle = 9°) with 1 mm sagittal slices. Functional echo planar images were acquired using a T2*-weighted sequence (TR = 1.0 s, TE = 30 ms, 44 slices, 2.0 mm thickness, in-plane resolution = 2 mm × 2 mm with 6/8 partial Fourier encoding, FOV = 212 mm, flip angle = 62°, multiband acceleration factor = 4); we included eight dummy volumes at the start of each scan to allow the scanner to reach B1 equilibrium. Functional images were acquired with anterior-to-posterior phase encoding; following each task, we collected two additional volumes with reversed phase encoding to correct for image distortion caused by B0 inhomogeneities.

The scanning session for *Alien Identity+Size* participants started with MPRAGE acquisition. Next, participants completed four runs of the tonotopy task (4 sweeps/run; 1^st^ and 3^rd^ up-sweep, 2^nd^ and 4^th^ down-sweep) during which they fixated on a central cross overlaid on a small landscape image to minimize eye movements associated with the acoustic sweep; images changed over the course each run (5 images/run; random duration to avoid aliasing with the acoustic sweep rate).

Next, there were six runs (10 blocks each, 8 trials/block) of 2AFC categorization. Across runs, 20 blocks involved categorization of alien identity across the high diagnostic frequency band, 20 involved identity categorization across the low frequency band, and 20 involved size categorization across amplitude. Use of the 2AFC (rather than 4AFC) categorization task in the scanner afforded several benefits: a) increasing power to differentiate across higher and lower frequency bands by minimizing switches within a block; b) reducing saccades across distinct visual aliens; c) constraining each block to two categories making identity and size tasks completely parallel. Our prior behavioral study demonstrated no learning advantage for even explicit instruction indicating the diagnostic frequency band (Obasih et al., 2023). A pseudorandomized block order common across participants assured that blocks reliant on the same diagnostic dimension did not occur consecutively. Overall block order in runs 1-3 repeated in runs 4-6, but with trial order shuffled, and with the amplitude set to a different level within the ‘big’ or ‘small’ amplitude range. In all, there were 320 alien identity categorization trials (80/category, with equal exemplars from the two size categories) and 160 alien size categorization trials (with equal exemplars from the four identity categories).

To facilitate comparisons between the alien identity and alien size categorization blocks, 20 of the 40 blocks that appeared during the alien identity task also appeared in the alien size task, with the same fixed trial order within the block. Half of the alien size blocks appeared before their corresponding alien identity blocks, and half appeared after. Each block began with a 3-sec visual display of the task (“Alien Identity” or “Alien Size”) followed by 1.5 sec for the audio and a 1.5 sec response window.

Finally, there were two runs of the attention-o-tonotopy task (one with base frequencies of 280 and 1253 Hz, one with base frequencies of 400 and 2123 Hz). Each run involved 15 blocks (22-34 sequence/block), with 12-15 sec of rest after every 3 blocks (except after the final block in a run). Auditory instructions at the beginning of each block indicated the locus of attention (“high,” “low,” “volume,” “rest”). As in the tonotopy task, participants fixated on a central cross overlaid on changing landscape images to avoid high- or low-band-attention-related eye movement (5 images/run).

The *Alien Size* participant group completed an abbreviated MRI protocol that included structural imaging, four runs of the tonotopy task, and two runs of 2AFC categorization. Critically, the 2AFC categorization task only involved alien size trials that followed the same alien size trials (in the same order) as the *Alien Identity+Size* group.

#### MRI Analyses

Cortical surfaces were reconstructed for each participant from the T1-weighted MPRAGE using FreeSurfer (Dale & Sereno, 1993; Dale, Fischl, & Sereno, 1999; Fischl, 2012).

Functional images from the tonotopy task were minimally preprocessed using AFNI (Cox, 1996) to account for saturation by discarding the first 8 volumes, to unwarp images using phase-reversed images collected at the end of each task, and to align all volumes to a reference volume from the middle of the first run. Next, the reference volume was aligned to the cortical surface in Freesurfer using boundary-based registration (Greve & Fischl, 2009).

To establish the regions of auditory cortex showing frequency selectivity within and across subjects, we calculated tonotopic maps using Fourier-based analyses on the preprocessed functional data from the tonotopy task in csurf, following standard phase-encoded mapping approaches (Sereno et al., 1995; Dick et al. 2012, 2017). Relying on the fact that each frequency step occurs at a consistent time within a sweep, Fourier analysis allowed us to compute a set of F-statistics indicating how each voxel responds at the frequency of stimulus cycling (4 cycles per run, 4 64-sec sweeps per run) relative to other frequencies; the phase lag (i.e., the delay relative to the start of the cycle) of the maximal response can be used to determine the frequency preference of each voxel. We performed Fourier analysis of each run (up-sweeps time-reversed for phase averaging with down-sweeps).

Finally, we painted the phase of the signal (indicating frequency preference) as a color map and visualized the data on each participant’s cortical surface reconstruction. Projecting each subject’s data onto an icosahedral surface and averaging across participants created a group map that was projected to a single participant’s surface reconstruction for visualization. Subsequent region-of-interest (ROI) analyses, described below, were limited to regions in auditory cortex where tonotopic organization was observed at the group level in participants expert in alien-identity categorization.

We next performed a regression-based analysis on the preprocessed functional data from each of the three tasks using 3dREMLfit in AFNI (Chen et al., 2012). For each analysis, we constructed regressors by convolving a vector of the onset times for each frequency step with a square wave of the appropriate duration. Rigid-body motion parameters (movement in the x-, y- and z-axis directions, as well as pitch, roll, and yaw) were included as regressors of no interest. The resultant beta estimates were used to compute, for each task, the difference in activation observed according to higher versus lower frequency band.

In the categorization tasks, this corresponded to the difference in blocks involving categorization of alien identity across the higher frequency diagnostic band versus those involving the lower frequency diagnostic band. In the attention tonotopy task, this corresponded to the difference between explicitly attending to high-versus low-frequency mini-sequences. Results were mapped to the cortical surface using the mri_vol2surf Freesurfer command. To generate group-level contrast maps, subject-level maps were projected onto an icosahedral mesh and averaged, the result of which was displayed on a single subject’s surface.

Although there is a macroscopic cross-participant similarity in large-scale tonotopic gradients, there is considerable cross-participant variability in frequency preference at a finer grain, particularly when voxels are aligned by cortical folding patterns (Besle et al., 2018; Moerel et al., 2014). Thus, to measure regional encoding and cross-task concordance in frequency selectivity while taking this variability into account, we calculated the degree of concordance in cross-task voxel-wise frequency selectivity in a set of small cortical surface ROIs defined in a previous study (Dick et al., 2017, see **Fig 3**). Here, we limit our analysis to those ROIs where tonotopic selectivity was observed at the group level in our alien-identity categorization experts. For each participant, the ROIs that tessellate tonotopically organized auditory cortex (37 per hemisphere) were projected from a single participant to the unit icosahedron (sphere_reg), then projected again to each curvature-aligned participant’s surface, and finally to the bbregister-aligned native space 2 x 2 x 2 mm EPI space using Freesurfer’s mri_label2vol.

For each ROI, cross-task concordance was evaluated via regression analysis, where all subject-wise beta coefficients in one task (e.g., *b*_tonotopy(high-low)_) were used to predict the high-low betas in another task (e.g., *b*_attention-o-tonotopy(high-low)_); each regression model included random intercepts for each participant. The resultant *t* values were transformed to *z* statistics, and these *z* values were painted onto the cortical surface. Significance (alpha < 0.05) was assessed using one-tailed tests, as we had *a priori* hypothesized a positive relationship between betas across tasks. To control for multiple comparisons, we applied a false discovery rate (FDR) correction on our *p* values using the 3dFDR command in AFNI. Note that because 3dFDR performs a two-tailed adjustment, *p* values associated with negative *z* statistics were inverted (i.e., 1-p) both prior to and following FDR correction, yielding FDR-corrected values that reflect one-tailed tests. Similar analyses were performed for different subgroups of participants defined based on behavioral performance during training.

We also conducted a series of analyses to compare cortical responses between subgroups of *Alien Identity+Size* participants (e.g., in participants who acquired category expertise early in training vs. those who acquired expertise over the course of training; see **Fig 5**) as well as compared to *Alien Size* participants. We tested whether the resultant cross-task concordance statistics differed across groups by computing a z_difference_ statistic (Hays, 1994), defined as:

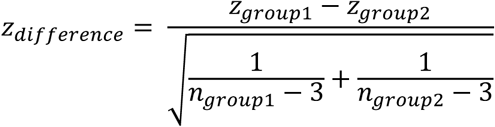

These values were projected onto a single subject’s cortical surface, providing an index of which group demonstrated stronger cross-task concordance. For these analyses, significance was assessed using two-tailed tests and FDR corrections were applied to control for multiple comparisons.

## Data Availability

The datasets and code generated and analyzed in the current study are available at https://app.box.com/folder/337623928007.

## Acknowledgments

The work was supported by grants from National Institutes of Health (R01DC004674) to LLH AT, and FD, the National Science Foundation (SBE/BCS 2414066, 2420979) to LLH and FD, an individual NRSA from the National Institute on Deafness and Communication Disorders to SL (F32DC020625) under the mentorship of LLH and FD. The authors would like to thank Jenny Bizley, Ayelet Landau, Rob Leech, Federico de Martino, Michelle Moerel, and Jeremy Skipper for comments on previous drafts of this manuscript.

## Author Contributions

**SL** contributed to Conceptualization, Methodology, Software, Formal Analysis, Investigation, Data Curation, Writing – Original Draft, Writing – Review and Editing, Visualization. **RR** contributed to Investigation, Data Curation, Writing – Original Draft. **ATT** contributed to Conceptualization, Methodology, Investigation, Funding Acquisition. **LLH** contributed to Conceptualization, Methodology, Investigation, Writing – Original Draft, Writing – Review and Editing, Visualization, Supervision, Resources, Project Administration, Funding Acquisition. **FD** contributed to Conceptualization, Methodology, Formal Analysis, Investigation, Writing – Original Draft, Writing – Review and Editing, Visualization, Supervision, Project Administration, Resources, Funding Acquisition.

## Materials and Correspondence

Correspondence and materials requests should be directed to Fred Dick.

## Supplemental Materials

**Supplemental Figure S1.**
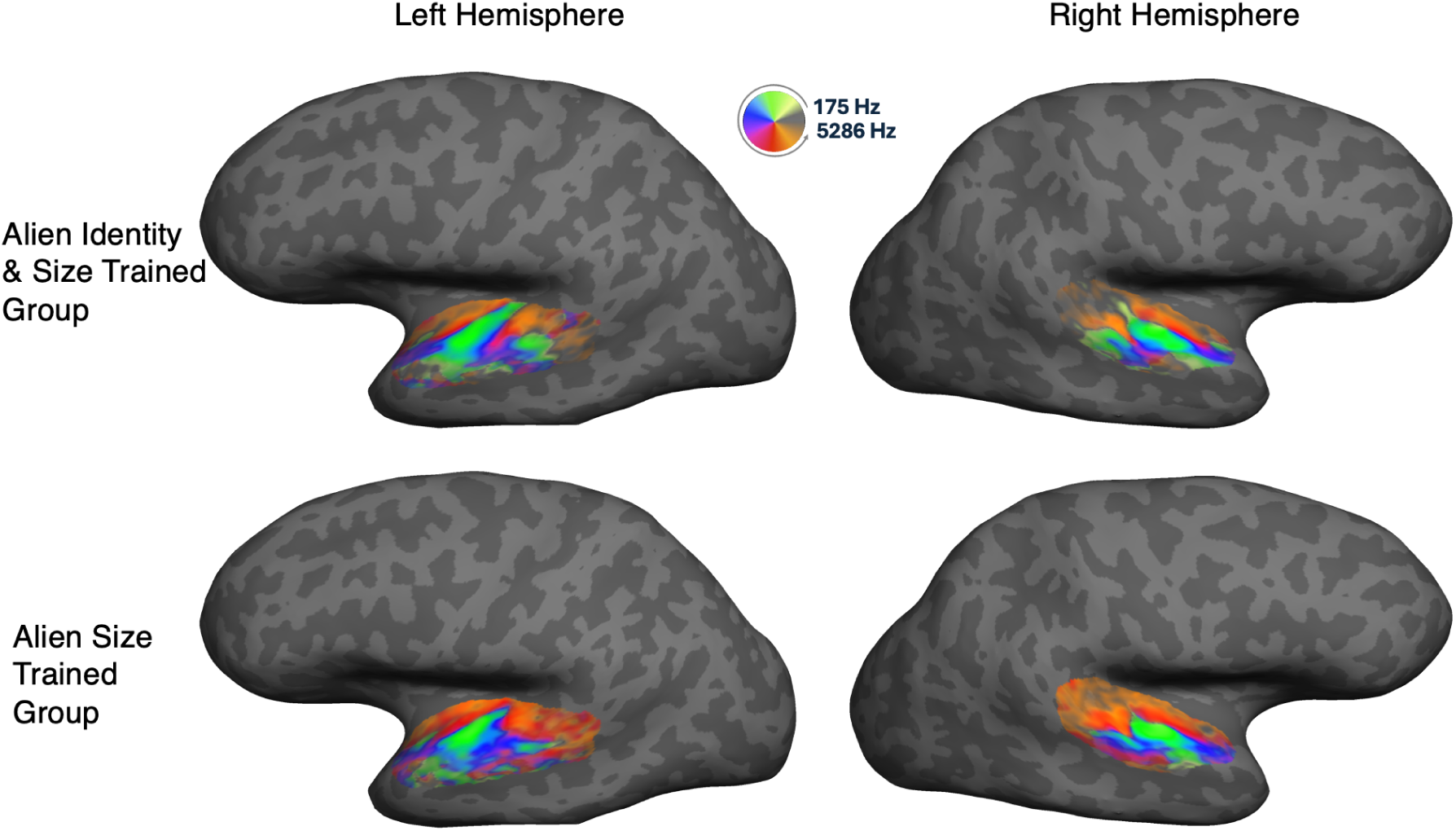
Cortical-surface-based group averages of phase-encoded tonotopic maps from the alien identity + size training group (top) and alien size training group (bottom). Color mapping shows phase of maximal fMRI response at 4 cycles/run; this indexes preferred auditory frequency via the high-to-low or low-to-high frequency sweep that repeats 4 times during each tonotopy run. (See Dick et al., 2012 and Methods for details).

**Supplemental Figure S2.**
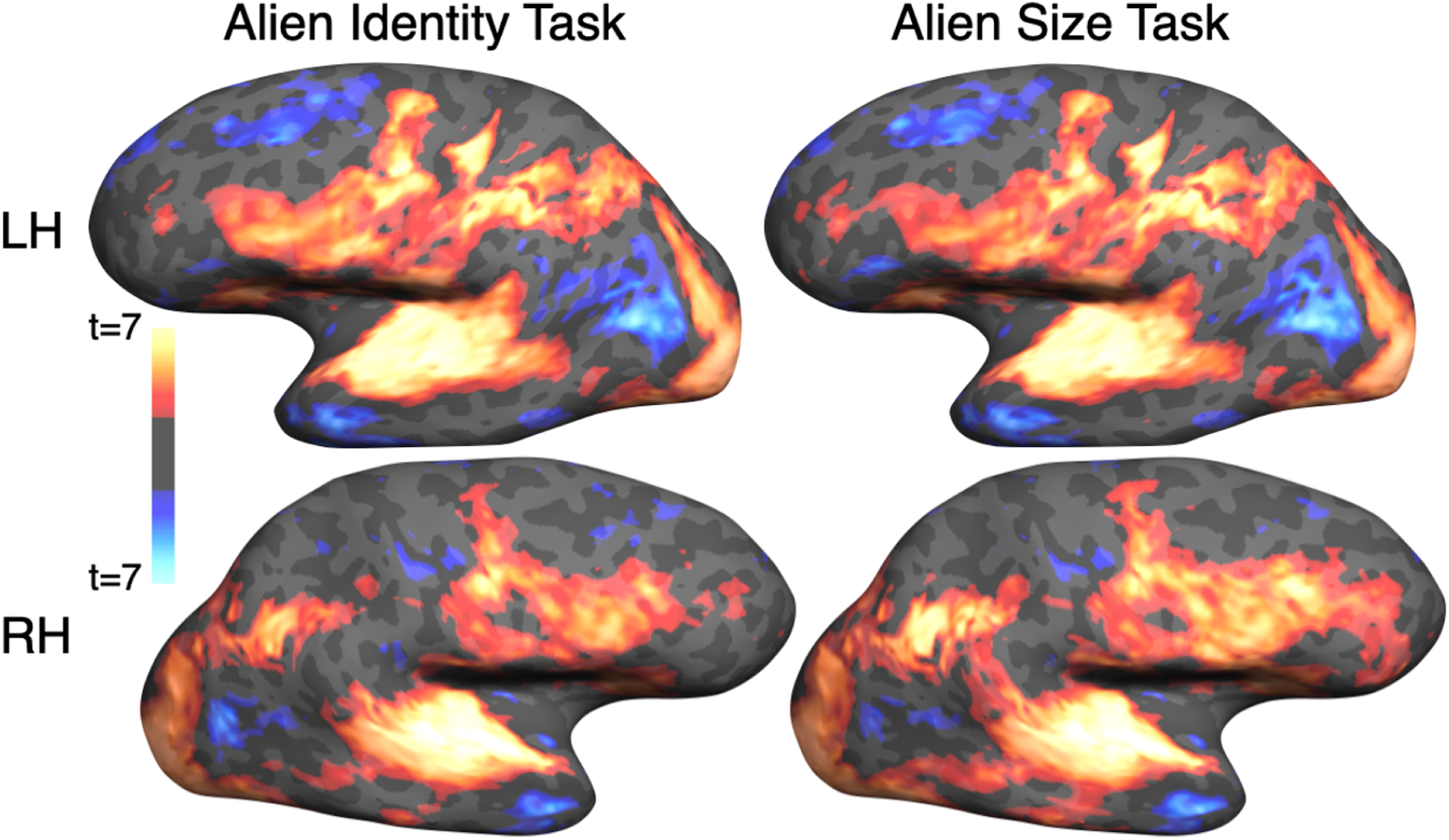
Cortical-surface-based group averages of BOLD response to the alien identity and alien size task compared to resting baseline, for experts trained on alien identity and alien size. Average brain-wise responses to both tasks with the auditory category exemplars show extensive bilateral activation of auditory-related cortex in temporal, inferior parietal, and frontal regions, as well as lateral occipital cortical regions typically associated with visual object recognition.

## References

Atiani, S., David, S. V., Elgueda, D., Locastro, M., Radtke-Schuller, S., Shamma, S. A., & Fritz, J. B. (2014). Emergent Selectivity for Task-Relevant Stimuli in Higher-Order Auditory Cortex. Neuron, 82(2), 486–499. 10.1016/j.neuron.2014.02.029

Bagur, S., Averseng, M., Elgueda, D., David, S., Fritz, J., Yin, P., Shamma, S., Boubenec, Y., & Ostojic, S. (2018). Go/No-Go task engagement enhances population representation of target stimuli in primary auditory cortex. Nature Communications, 9(1), 2529. 10.1038/s41467-018-04839-9

Besle, J., Mougin, O., Sánchez-Panchuelo, R.-M., Lanting, C., Gowland, P., Bowtell, R.,… Krumbholz, K. (2018). Is Human Auditory Cortex Organization Compatible With the Monkey Model? Contrary Evidence From Ultra-High-Field Functional and Structural MRI. Cerebral Cortex, 28(Suppl 1), 287–19. 10.1093/cercor/bhy267

Beeck, H. P. O. de, Pillet, I., & Ritchie, J. B. (2019). Factors Determining Where Category-Selective Areas Emerge in Visual Cortex. Trends in Cognitive Sciences, 23(9), 784–797. 10.1016/j.tics.2019.06.006

Bilalić, M. (2016). Revisiting the Role of the Fusiform Face Area in Expertise. Journal of Cognitive Neuroscience, 28(9), 1345–1357. 10.1162/jocn_a_00974

Chandrasekaran, B., Sampath, P. D., & Wong, P. (2010). Individual variability in cue-weighting and lexical tone learning. The Journal of the Acoustical Society of America, 128(1), 456–465.

Chua, K. W., Richler, J. J., & Gauthier, I. (2015). Holistic processing from learned attention to parts. Journal of Experimental Psychology: General, 144(4), 723–729. 10.1037/xge0000063

Collina, J. S., Erdil, G., Xia, M., Angeloni, C. F., Wood, K. C., Sheth, J., Kording, K. P., Cohen, Y. E., & Geffen, M. N. (2025). Individual-specific strategies inform category learning. Scientific reports, 15(1), 2984. 10.1038/s41598-024-82219-8

Da Costa, S., Zwaag, W. V. D., Miller, L. M., Clarke, S., & Saenz, M. (2013). Tuning in to sound: frequency-selective attentional filter in human primary auditory cortex. Journal of Neuroscience, 33(5), 1858–1863. 10.1523/jneurosci.4405-12.2013

Da Costa, S., van der Zwaag, W., Marques, J. P., Frackowiak, R. S., Clarke, S., & Saenz, M. (2011). Human primary auditory cortex follows the shape of Heschl’s gyrus. The Journal of Neuroscience, 31(40), 14067–14075. 10.1523/JNEUROSCI.2000-11.2011

De Baene, W., Ons, B., Wagemans, J., & Vogels, R. (2008). Effects of category learning on the stimulus selectivity of macaque inferior temporal neurons. Learning & Memory, 15(9), 717– 727. 10.1101/lm.1040508

DeGutis, J., & D’Esposito, M. (2007). Distinct mechanisms in visual category learning. Cognitive, affective & behavioral neuroscience, 7(3), 251–259. 10.3758/cabn.7.3.251

Deveau, J., Lovcik, G., & Seitz, A. R. (2014). Broad-based visual benefits from training with an integrated perceptual-learning video game. Vision Research, 99, 134–140. 10.1016/j.visres.2013.12.015

Dick, F. K., Lehet, M. I., Callaghan, M. F., Keller, T. A., Sereno, M. I., & Holt, L. L. (2017). Extensive Tonotopic Mapping across Auditory Cortex Is Recapitulated by Spectrally Directed Attention and Systematically Related to Cortical Myeloarchitecture. The Journal of neuroscience : the official journal of the Society for Neuroscience, 37(50), 12187–12201. 10.1523/JNEUROSCI.1436-17.2017

Emberson, L. L., Liu, R., & Zevin, J. D. (2013). Is statistical learning constrained by lower level perceptual organization? Cognition, 128(1), 82–102. 10.1016/j.cognition.2012.12.006

Folstein, J. R., Palmeri, T. J., & Gauthier, I. (2013). Category learning increases discriminability of relevant object dimensions in visual cortex. Cerebral Cortex (New York, N.Y.: 1991), 23(4), 814–823. 10.1093/cercor/bhs067

Folstein, J., Palmeri, T. J., Gulick, A. E. V., & Gauthier, I. (2015). Category Learning Stretches Neural Representations in Visual Cortex. Current Directions in Psychological Science, 24(1), 17–23. 10.1177/0963721414550707

Foster, J. J., & Ling, S. (2022). Feature-Based Attention Multiplicatively Scales the fMRI-BOLD Contrast-Response Function. The Journal of Neuroscience, 42(36), 6894–6906. 10.1523/JNEUROSCI.0513-22.2022

Freedman, D. J., Riesenhuber, M., Poggio, T., & Miller, E. K. (2003). A comparison of primate prefrontal and inferior temporal cortices during visual categorization. Journal of Neuroscience, 23, 5235–5246.

Gao, M., Turner, B. M., & Sloutsky, V. M. (2024). The Role of Attention in Category Representation. Cognitive Science, 48(4), e13438. 10.1111/cogs.13438

Gillebert, C. R., Op de Beeck, H. P., Panis, S., & Wagemans, J. (2008). Subordinate categorization enhances the neural selectivity in human object-selective cortex for fine shape differences. Journal of Cognitive Neuroscience, 21, 1054–1064.

Gomez, J., Barnett, M., & Grill-Spector, K. (2019). Extensive childhood experience with Pokémon suggests eccentricity drives organization of visual cortex. Nature Human Behaviour, 10.1038/s41562-019-0592-8

Gundlach, C., Wehle, S., & Müller, M. M. (2023). Early sensory gain control is dominated by obligatory and global feature-based attention in top-down shifts of combined spatial and feature-based attention. Cerebral cortex (New York, N.Y. : 1991), 33(19), 10286–10302. 10.1093/cercor/bhad282

Hays, W. L. (William L. (1994). Statistics / William L. Hays (5th ed). Harcourt Brace College Publishers.

Heald, S. L. M., & Nusbaum, H. C. (2014). Speech perception as an active cognitive process. Frontiers in Systems Neuroscience, 8, 35. 10.3389/fnsys.2014.00035

Haist, F., Lee, K., & Stiles, J. (2010). Individuating faces and common objects produces equal responses in putative face-processing areas in the ventral occipitotemporal cortex. Frontiers in Human Neuroscience, 4, 181. 10.3389/fnhum.2010.00181

Hausfeld, L., Riecke, L., & Formisano, E. (2018). Acoustic and higher-level representations of naturalistic auditory scenes in human auditory and frontal cortex. NeuroImage, 173, 472–483. 10.1016/j.neuroimage.2018.02.065

Heynckes, M., Lage-Castellanos, A., Weerd, P. D., Formisano, E., & Martino, F. D. (2023). Layer-specific correlates of detected and undetected auditory targets during attention. Current Research in Neurobiology, 4, 100075. 10.1016/j.crneur.2023.100075

Holmes, E., Domingo, Y., & Johnsrude, I. S. (2018). Familiar Voices Are More Intelligible, Even if They Are Not Recognized as Familiar. Psychological Science, 29(10), 1575–1583. 10.1177/0956797618779083

Idemaru, K., & Holt, L. L. (2013). The developmental trajectory of children’s perception and production of English /r/-/l/. The Journal of the Acoustical Society of America, 133(6), 4232– 4246. 10.1121/1.4802905

Jiang, X., Bradley, E., Rini, R. A., Zeffiro, T., Vanmeter, J., & Riesenhuber, M. (2007). Categorization training results in shape- and category-selective human neural plasticity. Neuron, 53, 891–903.

Jiang, X., Chevillet, M. A., Rauschecker, J. P., & Riesenhuber, M. (2018). Training Humans to Categorize Monkey Calls: Auditory Feature- and Category-Selective Neural Tuning Changes. Neuron, 98(2), 405–416.e4. 10.1016/j.neuron.2018.03.014

Karmiloff-Smith A. (2010). Neuroimaging of the developing brain: taking "developing" seriously. Human brain mapping, 31(6), 934–941. 10.1002/hbm.21074

Kraus, M. W., Torrez, B., Park, J. W., & Ghayebi, F. (2019). Evidence for the reproduction of social class in brief speech. Proceedings of the National Academy of Sciences of the United States of America, 116(46), 22998–23003. 10.1073/pnas.1900500116

Krumhuber, E.G., Skora, L.I., Hill, H.C.H. et al. The role of facial movements in emotion recognition. Nature Reviews Psycholology, 2, 283–296 (2023). 10.1038/s44159-023-00172-1

Kruschke, J. K., & Johansen, M. K. (1999). A model of probabilistic category learning. Journal of Experimental Psychology: Learning, Memory, and Cognition, 25(5), 1083.

Lavan N. (2022). The effect of familiarity on within-person age judgements from voices. British Journal of Psychology, 113(1), 287–299. 10.1111/bjop.12526

Leech, R., Holt, L. L., Devlin, J. T., & Dick, F. (2009). Expertise with artificial nonspeech sounds recruits speech-sensitive cortical regions. The Journal of neuroscience : the official journal of the Society for Neuroscience, 29(16), 5234–5239. 10.1523/JNEUROSCI.5758-08.2009

Ley, A., Vroomen, J., Hausfeld, L., Valente, G., De Weerd, P., & Formisano, E. (2012). Learning of new sound categories shapes neural response patterns in human auditory cortex. The Journal of Neuroscience, 32(38), 13273–13280. 10.1523/JNEUROSCI.0584-12.2012

Lim, S. J., Fiez, J. A., & Holt, L. L. (2019). Role of the striatum in incidental learning of sound categories. Proceedings of the National Academy of Sciences of the United States of America, 116(10), 4671–4680. 10.1073/pnas.1811992116

Liu, S. T., Montes-Lourido, P., Wang, X., & Sadagopan, S. (2019). Optimal features for auditory categorization. Nature Communications, 10(1), 1302. 10.1038/s41467-019-09115-y

Love, B. C., Medin, D. L., & Gureckis, T. M. (2004). SUSTAIN: A Network Model of Category Learning. Psychological Review, 111(2), 309–332. 10.1037/0033-295x.111.2.309

Martino, F. D., Moerel, M., Ugurbil, K., Goebel, R., Yacoub, E., & Formisano, E. (2015). Frequency preference and attention effects across cortical depths in the human primary auditory cortex. Proceedings of the National Academy of Sciences of the United States of America, 112(52), 16036–16041. 10.1073/pnas.1507552112

McColeman, C. M., Barnes, J. I., Chen, L., Meier, K. M., Walshe, R. C., & Blair, M. R. (2014). Learning-Induced Changes in Attentional Allocation during Categorization: A Sizable Catalog of Attention Change as Measured by Eye Movements. PLoS ONE, 9(1), e83302. 10.1371/journal.pone.0083302

McGugin, R. W., Gatenby, J. C., Gore, J. C., & Gauthier, I. (2012). High-resolution imaging of expertise reveals reliable object selectivity in the fusiform face area related to perceptual performance. Proceedings of the National Academy of Sciences of the United States of America. 10.1073/pnas.111633310

McKeeff, T. J., McGugin, R. W., Tong, F., & Gauthier, I. (2010). Expertise increases the functional overlap between face and object perception. Cognition, 117(3), 355–360. 10.1016/j.cognition.2010.09.002

McKone, E., Kanwisher, N., & Duchaine, B. C. (2007). Can generic expertise explain special processing for faces? Trends in Cognitive Sciences, 11(1), 8–15. 10.1016/j.tics.2006.11.002

McMurray, B., Danelz, A., Rigler, H., & Seedorff, M. (2018). Speech categorization develops slowly through adolescence. Developmental Psychology, 54(8), 1472–1491. 10.1037/dev0000542

Moerel, M., Martino, F. D., & Formisano, E. (2014). An anatomical and functional topography of human auditory cortical areas. Frontiers in Neuroscience, 8. Retrieved from 10.3389/fnins.2014.00225

Moerel, M., Yacoub, E., Gulban, O. F., Lage-Castellanos, A., & Martino, F. D. (2021). Using high spatial resolution fMRI to understand representation in the auditory network. Progress in Neurobiology, 207, 101887. 10.1016/j.pneurobio.2020.101887

Montes-Lourido, P., Kar, M., David, S. V., & Sadagopan, S. (2021). Neuronal selectivity to complex vocalization features emerges in the superficial layers of primary auditory cortex. PLoS Biology, 19(6), e3001299. 10.1371/journal.pbio.3001299

Murphy, T. K., Nozari, N., & Holt, L. L. (2024). Transfer of statistical learning from passive speech perception to speech production. Psychonomic Bulletin & Review, 31(3), 1193–1205. 10.3758/s13423-023-02399-8

Norman-Haignere, S., Kanwisher, N. G., & McDermott, J. H. (2015). Distinct Cortical Pathways for Music and Speech Revealed by Hypothesis-Free Voxel Decomposition. Neuron, 88(6), 1281–1296. 10.1016/j.neuron.2015.11.035

Nosofsky R. M. (1986). Attention, similarity, and the identification-categorization relationship. Journal of Experimental Psychology: General, 115(1), 39–61. 10.1037//0096-3445.115.1.39

O’Bryan, S. R., Jung, S., Mohan, A. J., & Scolari, M. (2024). Category Learning Selectively Enhances Representations of Boundary-Adjacent Exemplars in Early Visual Cortex. The Journal of Neuroscience, 44(3), e1039232023. 10.1523/JNEUROSCI.1039-23.2023

Obasih, C. O., Luthra, S., Dick, F., & Holt, L. L. (2023). Auditory category learning is robust across training regimes. Cognition, 237, 105467. 10.1016/j.cognition.2023.105467

Petkov, C. I., Kang, X., Alho, K., Bertrand, O., Yund, E. W., & Woods, D. L. (2004). Attentional modulation of human auditory cortex. Nature Neuroscience, 7(6), 658–663. 10.1038/nn1256

Reetzke, R., Xie, Z., Llanos, F., & Chandrasekaran, B. (2018). Tracing the Trajectory of Sensory Plasticity across Different Stages of Speech Learning in Adulthood. Current Biology, 28(9), 1419–1427.e4. 10.1016/j.cub.2018.03.026

Riecke, L., Peters, J. C., Valente, G., Kemper, V. G., Formisano, E., & Sorger, B. (2017). Frequency-Selective Attention in Auditory Scenes Recruits Frequency Representations Throughout Human Superior Temporal Cortex. Cerebral Cortex. 10.1093/cercor/bhw160

Roark, C. L., & Chandrasekaran, B. (2023). Stable, flexible, common, and distinct behaviors support rule-based and information-integration category learning. NPJ Science of Learning, 8(1), 14. 10.1038/s41539-023-00163-0

Roark, C. L., Lehet, M. I., Dick, F., & Holt, L. L. (2022). The Representational Glue for Incidental Category Learning Is Alignment With Task-Relevant Behavior. Journal of Experimental Psychology: Learning, Memory, and Cognition, 48(6), 769–784. 10.1037/xlm0001078

Roark, C. L., Paulon, G., Rebaudo, G., McHaney, J. R., Sarkar, A., & Chandrasekaran, B. (2024). Individual differences in working memory impact the trajectory of non-native speech category learning. PloS One, 19(6), e0297917. 10.1371/journal.pone.0297917

Roy, J. E., Riesenhuber, M., Poggio, T., & Miller, E. K. (2010). Prefrontal cortex activity during flexible categorization. The Journal of Neuroscience, 30(25), 8519–8528. 10.1523/JNEUROSCI.4837-09.2010

Roye, A., Schröger, E., Jacobsen, T., & Gruber, T. (2010). Is my mobile ringing? Evidence for rapid processing of a personally significant sound in humans. The Journal of Neuroscience : The Official Journal of the Society for Neuroscience, 30(21), 7310–7313. 10.1523/jneurosci.1113-10.2010

Russ, B. E., Lee, Y. S., & Cohen, Y. E. (2007). Neural and behavioral correlates of auditory categorization. Hearing Research, 229(1-2), 204–212. 10.1016/j.heares.2006.10.010

Saenz, M., Buracas, G. T., & Boynton, G. M. (2002). Global effects of feature-based attention in human visual cortex. Nature Neuroscience, 5(7), 631–632. 10.1038/nn876

Saffran, J. R., & Kirkham, N. Z. (2018). Infant Statistical Learning. Annual Review of Psychology, 69(1), 181–203. 10.1146/annurev-psych-122216-011805

Sasaki, Y., Nanez, J. E., & Watanabe, T. (2010). Advances in visual perceptual learning and plasticity. Nature Reviews Neuroscience, 11(1), 53–60. 10.1038/nrn2737

Serences, J. T., & Boynton, G. M. (2007). The representation of behavioral choice for motion in human visual cortex. The Journal of neuroscience : the official journal of the Society for Neuroscience, 27(47), 12893–12899. 10.1523/JNEUROSCI.4021-07.2007

Serre, T., Wolf, L., Bileschi, S., Riesenhuber, M., & Poggio, T. (2007). Robust object recognition with cortex-like mechanisms. IEEE transactions on pattern analysis and machine intelligence, 29(3), 411–426. 10.1109/TPAMI.2007.56

Sheth, J., Collina, J. S., Piasini, E., Kording, K. P., Cohen, Y. E., & Geffen, M. N. (2025). The interplay of uncertainty, relevance and learning influences auditory categorization. Scientific reports, 15(1), 3348. 10.1038/s41598-025-86856-5

Sigala, N., & Logothetis, N. K. (2002). Visual categorization shapes feature selectivity in the primate temporal cortex. Nature, 415(6869), 318–320. 10.1038/415318a

Sigala, N., & Logothetis, N. K. (2002). Visual categorization shapes feature selectivity in the primate temporal cortex. Nature, 415(6869), 318–320. 10.1038/415318

Todorov, A., Pakrashi, M., & Oosterhof, N. (2009). Evaluating faces on trustworthiness after minimal time exposure. Social Cognition, 27, 813–833. 10.1521/soco.2009.27.6.813

Treue, S., & Maunsell, J. H. (1999). Effects of attention on the processing of motion in macaque middle temporal and medial superior temporal visual cortical areas. The Journal of neuroscience : the official journal of the Society for Neuroscience, 19(17), 7591–7602. 10.1523/JNEUROSCI.19-17-07591.1999

van der Linden, M., van Ruennout, M., & Idefrey, P. (2010). Formation of category representations in superior temporal sulcus. Journal of Cognitive Neuroscience, 22, 1270– 1282.

Van Gulick, A. E., & Gauthier, I. (2014). The perceptual effects of learning object categories that predict perceptual goals. Journal of Experimental Psychology: Learning, Memory, and Cognition, 40(5), 1307–1320. 10.1037/a0036822

Wade, T., & Holt, L. L. (2005). Incidental categorization of spectrally complex non-invariant auditory stimuli in a computer game task. The Journal of the Acoustical Society of America, 118(4), 2618–2633. http://pubmed.gov/16266182

Whitton, J. P., Hancock, K. E., Shannon, J. M., & Polley, D. B. (2017). Audiomotor Perceptual Training Enhances Speech Intelligibility in Background Noise. Current Biology : CB, 27(21), 3237–3247.e6. 10.1016/j.cub.2017.09.014

Wikman, P., Rinne, T., & Petkov, C. I. (2019). Reward cues readily direct monkeys’ auditory performance resulting in broad auditory cortex modulation and interaction with sites along cholinergic and dopaminergic pathways. Scientific Reports, 9(1), 3055. 10.1038/s41598-019-38833-y

Yin, P., Strait, D. L., Radtke-Schuller, S., Fritz, J. B., & Shamma, S. A. (2020). Dynamics and Hierarchical Encoding of Non-compact Acoustic Categories in Auditory and Frontal Cortex. Current Biology : CB, 30(9), 1649–1663.e5. 10.1016/j.cub.2020.02.047

Yoo, S. A., Martinez-Trujillo, J. C., Treue, S., Tsotsos, J. K., & Fallah, M. (2022). Attention to visual motion suppresses neuronal and behavioral sensitivity in nearby feature space. BMC Biology, 20(1), 220. 10.1186/s12915-022-01428-7

Young, A. W., Frühholz, S., & Schweinberger, S. R. (2020). Face and Voice Perception: Understanding Commonalities and Differences. Trends in Cognitive Sciences, 24(5), 398–410. 10.1016/j.tics.2020.02.001

Yurovsky, D., & Frank, M. C. (2017). Beyond naïve cue combination: salience and social cues in early word learning. Developmental Science, 20(2), 10.1111/desc.12349. https://doi.org/10.1111/desc.12349

Yurovsky, D., Boyer, T. W., Smith, L. B., & Yu, C. (2013). Probabilistic cue combination: less is more. Developmental Science, 16(2), 149–158. 10.1111/desc.12011

Zhong, L., Zhang, Y., Duan, C. A., Deng, J., Pan, J., & Xu, N. L. (2019). Causal contributions of parietal cortex to perceptual decision-making during stimulus categorization. Nature Neuroscience, 22(6), 963–973. 10.1038/s41593-019-0383-6

